# Marginated aberrant red blood cells induce pathologic vascular stress fluctuations in a computational model of hematologic disorders

**DOI:** 10.1101/2023.05.16.541016

**Authors:** Xiaopo Cheng, Christina Caruso, Wilbur A. Lam, Michael D. Graham

## Abstract

Red blood cell (RBC) disorders affect billions worldwide. While alterations in the physical properties of aberrant RBCs and associated hemodynamic changes are readily observed, in conditions such as sickle cell disease and iron deficiency, RBC disorders can also be associated with vascular dysfunction. The mechanisms of vasculopathy in those diseases remain unclear and scant research has explored whether biophysical alterations of RBCs can directly affect vascular function. Here we hypothesize that the purely physical interactions between aberrant RBCs and endothelial cells, due to the margination of stiff aberrant RBCs, play a key role in this phenomenon for a range of disorders. This hypothesis is tested by direct simulations of a cellular scale computational model of blood flow in sickle cell disease, iron deficiency anemia, COVID-19, and spherocytosis. We characterize cell distributions for normal and aberrant RBC mixtures in straight and curved tubes, the latter to address issues of geometric complexity that arise in the microcirculation. In all cases aberrant RBCs strongly localize near the vessel walls (margination) due to contrasts in cell size, shape, and deformability from the normal cells. In the curved channel, the distribution of marginated cells is very heterogeneous, indicating a key role for vascular geometry. Finally, we characterize the shear stresses on the vessel walls; consistent with our hypothesis, the marginated aberrant cells generate large transient stress fluctuations due to the high velocity gradients induced by their near-wall motions. The anomalous stress fluctuations experienced by endothelial cells may be responsible for the observed vascular inflammation.

**Significance Statement:** A common and potentially life-threatening complication of blood cell disorders is inflammation and dysfunction of the vascular wall, for reasons that remain unclear. To address this issue, we explore a purely biophysical hypothesis involving red blood cells using detailed computational simulations. Our results show that red blood cells that are pathologically altered in cell shape, size, and stiffness, which occurs in various blood disorders, strongly marginate, residing primarily in the cell-free layer near blood vessel walls, generating large shear stress fluctuations at the vessel wall that may be responsible for endothelial damage and inflammation.

---

**E**ndothelial cells lining the blood vessels of individuals with blood cell disorders are often dysfunctional and in a pro-inflammatory state, increasing the risk of stroke and atherosclerosis (1–5). In particular, stroke, a significant cause of mortality in sickle cell disease (SCD), often occurs in highly tortuous cerebral arteries and is associated with endothelial inflammation and chronic vasculopathy. In patients with cardiovascular disease and iron deficiency anemia (IDA), improved disease outcomes were observed with iron supplementation and subsequent resolution of IDA.(6); however, the underlying pathophysiologic basis for the association remains unknown. The interplay among adhesive RBC-endothelial interactions, inflammatory cytokines, and hemolysis all contribute to vasculopathy in blood disorders, however the potential contribution of the altered physical properties of aberrant RBCs, particularly shape and stiffness, to the hemodynamic environment experienced by the vascular endothelium remains poorly understood. This topic is the focus of the present work.

Vascular geometries contribute to vasculopathy in blood disorders (7). The vascular system is comprised of diverse geometries, including normal complexities such as curves and bifurcations as well as pathologic ones such as aneurysms and stenoses, and variations in vascular geometry cause significant changes in the local shear stress profile during blood flow, which are known to induce endothelial proinflammatory responses (2, 4, 7, 8). Leveraging an endothelialized microfluidic model of multiple geometries, Mannino et al. (9) found that VCAM-1 and E-selectin expression, biomarkers of endothelial cell dysfunction, significantly correlated with shear stress variation and were most pronounced near bifurcation points. Furthermore, they found that endothelial cells exposed to SCD RBCs exhibited increased endothelial inflammation along the outside wall of the bend in the curved regions of vessels (10). These observations indicate that it is essential to understand the role of vascular geometric complexity on endothelial dysfunction in blood disorders.

Aberrant RBCs arising in blood disorders often have very different physical properties compared to healthy RBCs. A typical example is SCD, in which abnormal sickle hemoglobin polymerizes within RBCs upon deoxygenation, creating long fibers that pathologically disrupt cellular architecture(11), leading to increased membrane stiffness as well as loss of cellular volume secondary via dehydration. Subsequently, sickle RBCs are biophysically less deformable than normal cells and some subpopulations are distorted irreversibly into a sickle-like shape. Similarly, in samples of blood from IDA patients, Caruso et al.(12) identified a subpopulation of very small and poorly deformable iron deficiency RBCs (idRBCs). When exposed to COVID-19, morphologically normal RBCs exhibited a conformational change to sphero-echinocytes with reduced size and deformability (13). Relatedly, plasma from adult COVID-19 patients causes significant RBC aggregation under flow, and fibrinogen-mediated aggregation directly damages the endothelial glycocalyx (14). In hereditary spherocytosis (HS), genetic mutations affect RBC membrane proteins, breaking the linkage between the membrane skeleton and the lipid bilayer, causing membrane loss (15). As a result, instead of being biconcave discoids, RBCs become inflexible spherical cells called spherocytes.

The spatial distribution of the different cellular components of blood is nontrivial and depends on the relative physical properties of the different components. Normal RBCs migrate toward the center of a blood vessel, leaving an RBC-depleted cell-free layer (CFL) near vessel walls. In contrast, white blood cells (WBCs) and platelets tend to reside in these layers, a flow-induced segregation phenomenon called margination (16–18).

The segregation behavior during blood flow is substantially dictated by the contrasts in the cellular properties, such as shape, size, and deformability, of the various components. Kumar et al. (19, 20) used detailed simulations to probe the effect of rigidity difference in a binary suspension of deformable capsules in shear flow. They found that stiff capsules display substantial margination when they are the dilute component, while flexible capsules tend to enrich around the channel’s centerline. Similarly, in a mixture of large and small capsules,the smaller capsules marginate (20). Sinha et al. (21) investigated the flow-induced segregation behavior in binary suspensions of spherical and ellipsoidal capsules in simple shear flow by varying the aspect ratio while keeping constant either the equatorial radius or volume of capsules. Direct simulations with models of blood corroborate these model results (22, 23). A simple theory of margination based on the two key transport mechanisms of cells in flow – cell-cell collisions and hydrodynamic migration of deformable cells away from walls (24, 25) predicts that a subpopulation of rigid particles in a suspension of primarily deformable particles, will strongly concentrate at walls during flow (26, 27).

Margination may have particular significance in the context of vasculopathy in blood cell disorders. Endothelial cells are responsible for translating biophysical cues, such as the shear force of the hemodynamic microenvironment, into cellular biological signals (5, 28, 29). Pathological alterations of such forces promote endothelial activation with the release of proinflammatory signals (30–32), which contribute to atherosclerotic plaques susceptible to myocardial infarction and strokes(33). Indeed, the fact that vasculopathy pervasively occurs even in the oxygenated conditions in both small and larger vessels demands a new understanding of SCD pathophysiology in the absence of vaso-occlusion, which occurs only under the deoxygenated conditions in the microvessels.

Inspired by advances in the mechanistic understanding of the distribution and segregation behaviors during blood flow as well as experimental observations of blood disorders during flow (9, 10, 12, 34), we propose a biophysical hypothesis for the pathophysiology of vasculopathy in blood disorders: diseased cells strongly marginate, residing primarily in the CFL near the vascular walls, resulting in endothelial inflammation by provoking fluctuations in local wall shear stress, which is consistent with the chronic and diffusive nature of vasculopathy in blood cell disorders. Limited computational studies of this hypothesis, for blood flow in straight tubes, have been performed for the cases of SCD (35) and IDA (12).

The present work uses detailed simulations of a cellular-scale mathematical model of blood flow in small vessels to examine this hypothesis. Several diseases are considered: SCD, IDA, COVID-19, and spherocytosis, in both a simple cylindrical blood vessel geometry and a more geometrically complex serpentine curved tube. The choice of these disease models arose from a number of considerations. On biophysical grounds, all of these disorders result in subpopulations of red cells with altered physical, morphological, and geometric properties. Biologically, they represent a spectrum of disorders that encompass different etiologies, illustrating the generalizability of our findings: SCD arises from a genetic disorder, iron deficiency anemia a nutritional one; COVID-19 is an example of an infectious disease that gives rise to biophysically altered red cell subpopulations, and spherocytosis can arise in genetic or acquired disorders. Not only do the results provide strong and broad-based computational support for our hypothesis, but they also begin to reveal transient aspects of the stress environment experienced by endothelial cells as well as the strong spatial variations in this environment engendered by a complex flow geometry.

## Results and discussion

### Model summary

We simulate a flowing suspension of RBCs, modeled as deformable fluid-filled elastic capsules, in rigid straight (Fig.1) and curved (Fig.3) cylindrical tubes with diameter *D* = 40*μm*. For the blood disease cases, RBC suspensions are modeled as binary mixtures of normal RBCs with aberrant RBCs from different blood disorders (e.g., idRBCs, sickle RBCs, sphero-echinocytes, and spherocytes). In the binary suspensions, the number fraction for normal RBCs is 0.9, and for aberrant RBCs is 0.1. This is a simplification, as in any real blood cell population there will be a distribution of cell properties. A suspension of only normal RBCs, referred to as healthy RBC suspension, is considered as a control. The overall volume fraction (tube hematocrit) is around 20%, consistent with the observed substantial decrease of hematocrit from large vessel to the microcirculation (37, 38). (There is some variation between the cases we consider here because different cell types have different volumes – what we keep constant between cases are number fraction and number density.) The suspending fluid, blood plasma, is considered incompressible and Newtonian with a viscosity of about *η* = 1.10 −1.35mPas. The discoid radius *a* for human RBC is about 4*μm*. The RBC membrane in-plane shear elasticity modulus *G* ∼ 2.5− 6*μ*N/m. The deformability of a capsule in the pressure-driven flow is measured by the dimensionless capillary number 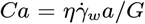 . *Ca* is set to be 1.0 for normal RBCs, which corresponds to 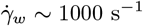. Since aberrant RBCs in blood disorders are generally much smaller and stiffer than normal RBCs, the interfacial shear modulus *G* of the aberrant RBCs is assumed to be five times that of normal RBCs, which leads to *Ca* for aberrant RBCs being at most 0.2 times that of *Ca* for normal RBCs. All results are for simulations that have been run to a statistically stationary state.

**Fig. 1.**
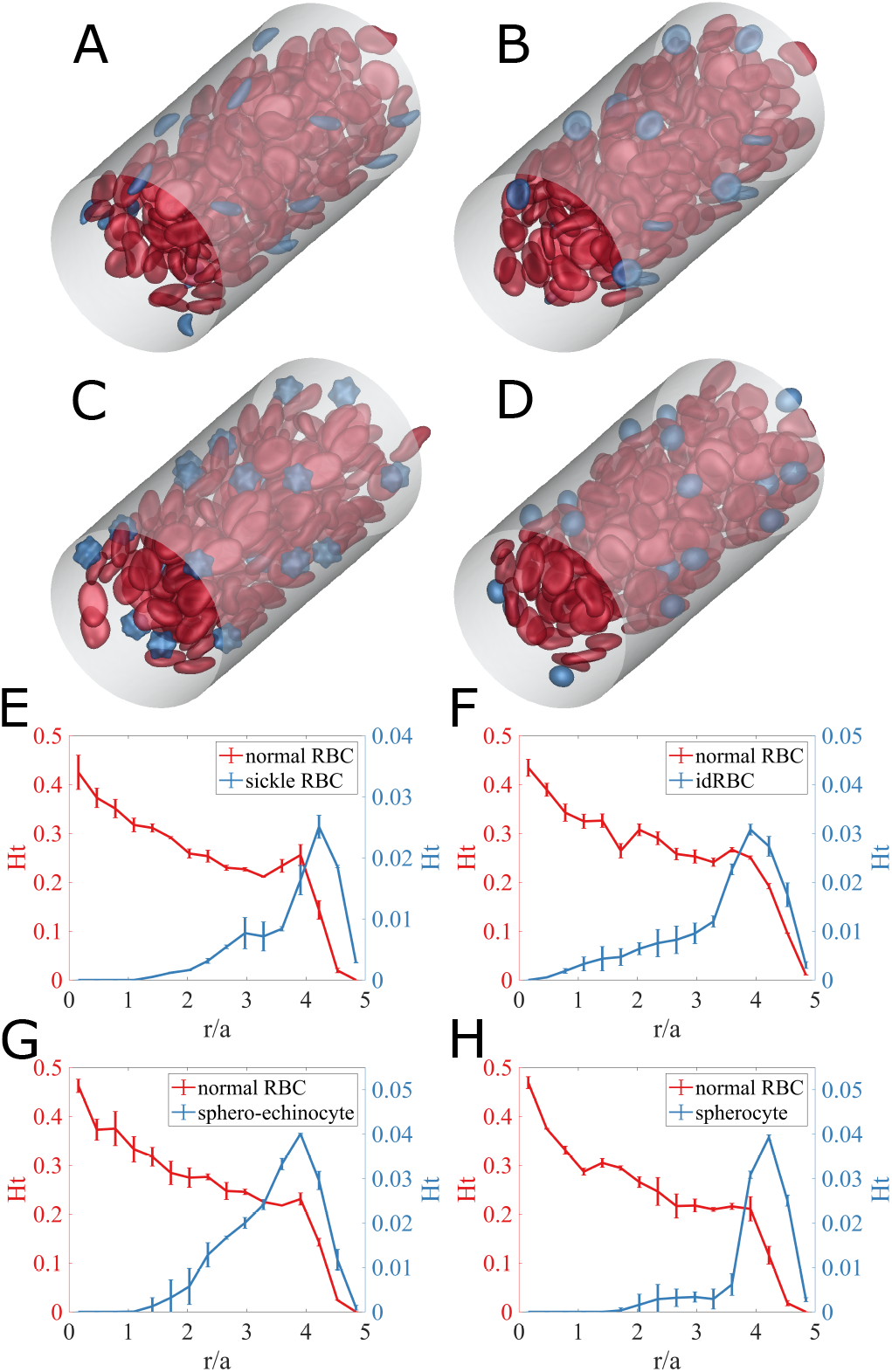
(Top) Simulation snapshots for (A) sickle cell disease, (B) iron deficiency anemia, (C) COVID-19, and (D) spherocytosis, in straight cylindrical tube subjected to a unidirectional pressure-driven flow. Red capsules represent normal RBCs with oblate spheroid shape for bending elasticity and biconcave discoid shape for shear elasticity, while blue capsules are for aberrant RBCs. (Bottom) Radial hematocrit profile for (red) normal RBCs and (blue) aberrant RBCs of (E) sickle cells in SCD, (F) idRBCs in IDA, (G) sphero-echinocytes in COVID-19, and (H) spherocytes in spherocytosis. The error bars represent estimated standard error using the block averaging method (36).

The spontaneous shape of the RBC membrane is inhomogeneous. Dupire et al. (39) showed that an RBC maintains its biconcave shape even during tank-treading and hypothesizes that this effect might come from anisotropic elastic properties or an inhomogeneous natural shape. Fischer et al.(40) found that RBCs have “shape memory”, which arises from spatial variations in their natural shape. The choice of the spontaneous shape can strongly affect the stable dynamics of the RBC. Sinha et al.(41) investigate the cell dynamics’ dependence on the membrane’s spontaneous curvature. They found that an oblate spheroidal spontaneous curvature maintains the dimple of the RBC during tank-treading dynamics and exhibits off-shear-plane, tumbling consistent with the experimental observations of Dupire et al.(39). For a complex structure such as an RBC membrane, it is possible that the natural shape for shear elasticity may differ from that for bending elasticity so the overall natural shape of an element results from the balance of bending and shear forces. Thus in this work, the spontaneous shape of RBC bending elasticity is taken to be the oblate spheroid, while the spontaneous shape of RBC shear elasticity is assumed to be the biconcave discoid. Further details are included in the Materials and Methods section and Supplementary Information; in particular we show that the results here are robust against changes in the details of the cell elasticity model.

### Cylindrical blood vessel

Figures 1(A-D) show snapshots from simulations of blood flow in a straight tube for SCD, IDA, COVID-19, and spherocytosis, respectively. (The SI contains movies of these simulations.) In all cases, the aberrant cells (blue) appear to be marginated. Fig.1(E-H), respectively, show the corresponding radial hematocrit profiles. These indicate that aberrant RBCs strongly marginate, while the normal RBCs display the expected CFL, and a concentration that increases toward the centerline. Sample simulations with doubled tube length and the same mesh spacing were also conducted; changes in the results were negligible. These results demonstrate that differences in cell size and deformability of the aberrant cells are sufficient to drive strong segregation behavior.

The presence of stiff and/or small aberrant cells near vessel walls is expected to generate high velocity gradients, and consequently large shear stresses on the walls, *τ*_*w*_. Fig.2(*A, B, C, D*) shows snapshots of the spatial distribution of excess wall shear stress 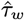 for the four cases. Here 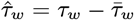 is defined as deviation from the mean wall shear stress 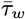. The red regions indicate large local fluctuations, and one can see that these are directly associated with nearby aberrant RBCs. Fig.2(*E*) shows time series of additional wall shear stress 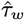 at a point on the wall for the various cases. Peaks of high additional wall shear stress are larger and more frequent in all of the disease cases than in the healthy case. These differences are further quantified in Fig.2(*F*), which shows the probability density profiles of excess wall shear stress in the suspensions. The PDFs for all aberrant RBC cases display a long tail at high 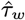, where the probability density of high wall shear stress for cases with aberrant RBCs is orders of magnitude higher than for the healthy case. This phenomenon is especially prominent for sphero-echinocytes and sickle RBCs and less pronounced with spherocytes, and is related to their morphology: the spiked surfaces of sphero-echinocytes and sickle RBCs induce high local wall shear stress, while the round spherocytes, though near the wall, roll smoothly without generating substantial excess stress.

**Fig. 2.**
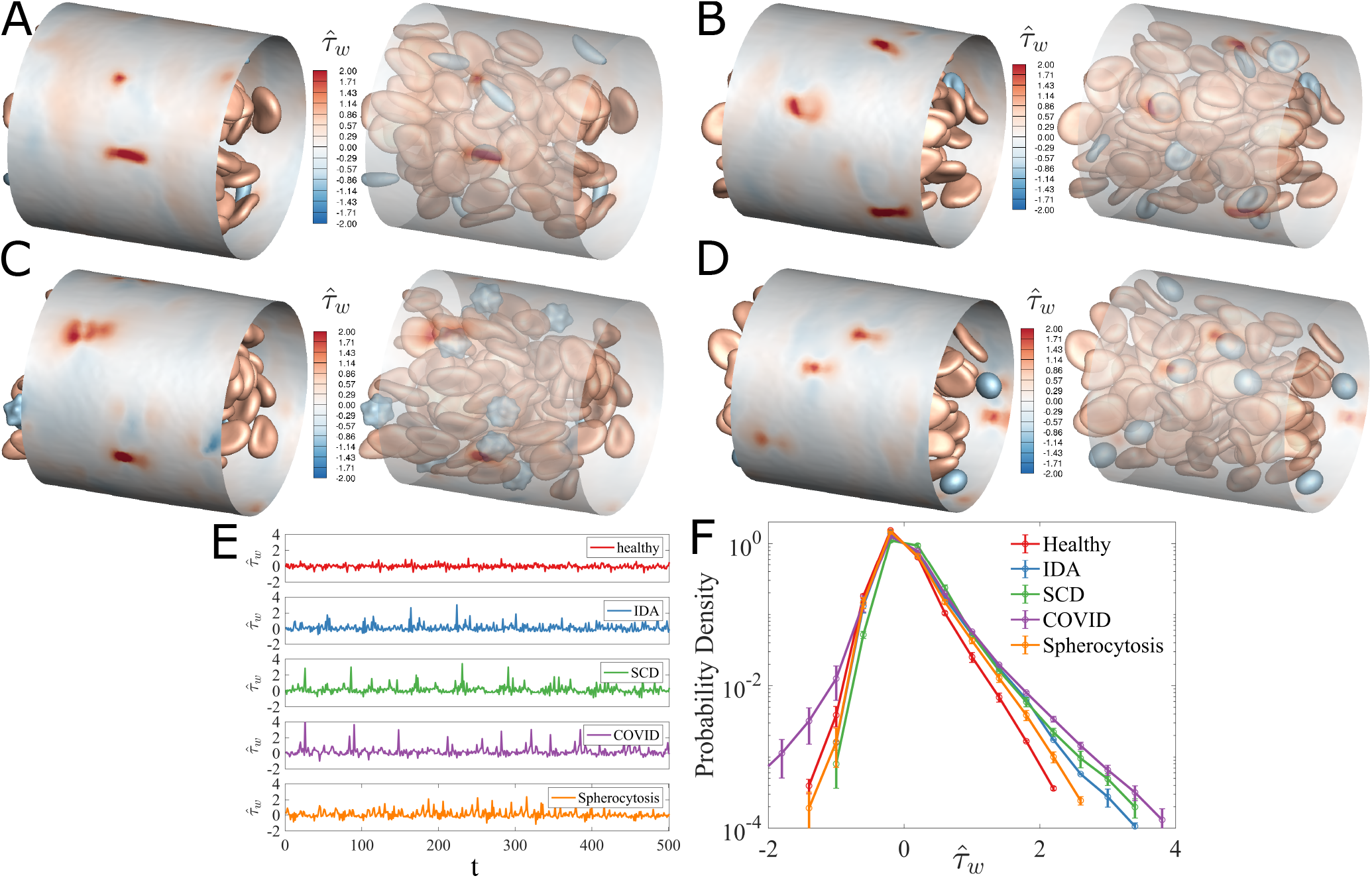
(Top) Simulation snapshots and corresponding transparent views of excess wall shear stress 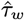, induced by the presence of the cells in (A) sickle cell diseases, (B) iron deficiency anemia, (C) COVID-19, and (D) spherocytosis RBC suspensions. To distinguish the colors of the cells themselves from the colors of the RBC-induced wall shear stress on the cylindrical surface, the color of normal RBCs is set to be pale red, and aberrant RBCs pale blue. (Bottom) (E) Time evolution of the additional wall shear stress 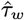 at a fixed wall position for the cases of a homogeneous suspension of healthy RBCs and binary suspensions of normal RBCs with aberrant RBCs, respectively. (F) Probability distribution of the additional wall shear stresses 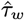 over the cylindrical wall in the various cases.

**Fig. 3.**
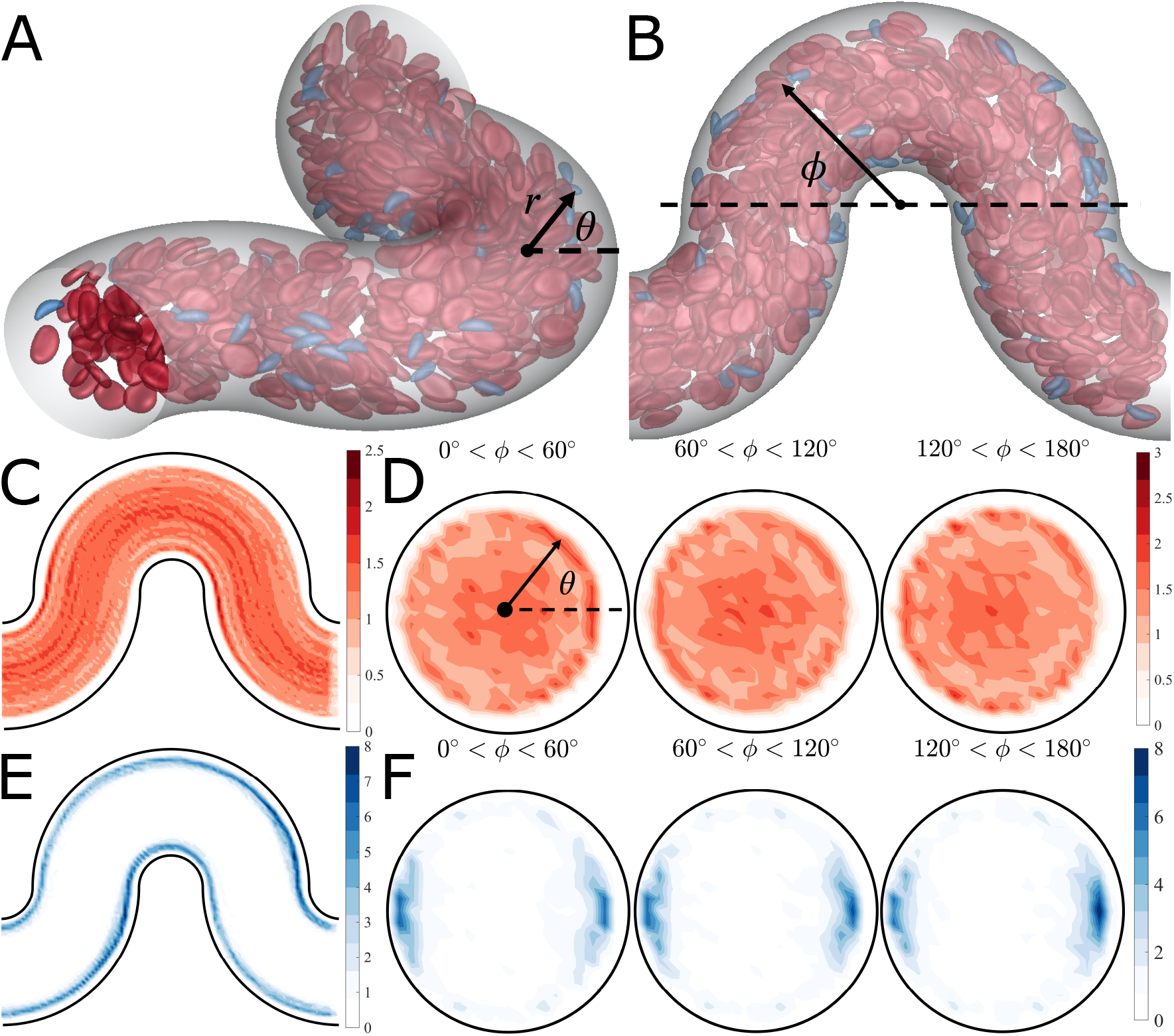
(A, B) Simulation snapshots of a suspension of normal RBCs with sickle RBCs in a curved channel, showing coordinate definitions for *r* the distance from the local channel centerline, *θ*, the relative position between the “inner side” (*θ* = 180°) and “outer side(*θ* = 0°)”, and *ϕ*, the angle around the curve. (C) Normalized center-plane cell number density and (D) normalized cross-sectional cell number density for normal RBCs in the binary SCD RBC suspension. (E) Normalized center-plane cell number density and (F) normalized cross-sectional cell number density for sickle RBCs in the binary SCD RBC suspension. Note that the curved channel is divided into three parts based on the value of angle *ϕ*; thus, the cross-sectional cell number distribution is computed over each part. The cell number density distributions are normalized so that if the spatial distributions of cells is uniform, then the normalized cell number density is unity everywhere within the curved channel.

### Curved blood vessel

We now consider cell and stress distributions in a curved tube. Fig.3, presents simulation results for a suspension of normal RBCs with sickle RBCs. Snapshots of cell distributions are shown in Fig.3(*A, B*), along with a coordinate system we use for the analysis. Fig.3 (*C, E*), show cell number density distributions for the (C) normal and (E) aberrant cells on the center plane of the channel. The margination of the aberrant cells is apparent. Fig.3(*D, F*) show the number density distributions for the normal and aberrant cells averaged over various segments of the channel, including both the normalized center-plane and cross-sectional cell distributions. Fig.3(*D*) shows that the CFL thickness is larger near the outer side (*θ* = 0°) and thinnest near the inner side (*θ* = 180°). Fig.3(*F*) indicates that sickle RBCs strongly focus at two near-wall locations, both at the inner and outer sides, on the centerplane. As the angle *ϕ* increases (i.e. as we move downstream around a bend), the concentration of sickle cells near the outer wall becomes more pronounced. Similar results are found for the other aberrant cell suspensions as well, as seen in the cross-sectional and centerplane distributions shown in Fig.4.

**Fig. 4.**
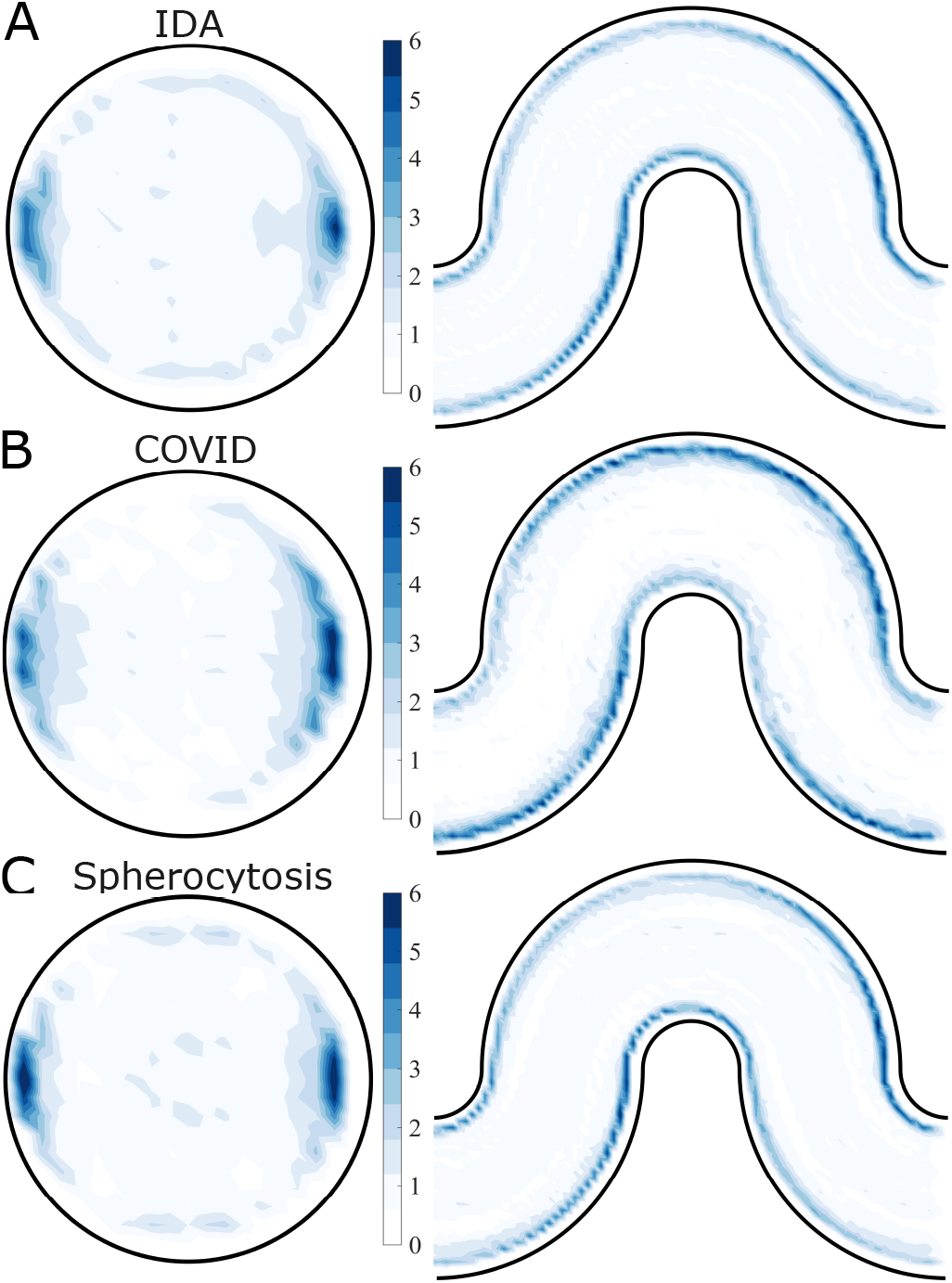
Cross-sectional and center-plane normalized cell number density distribution of (A) iron deficiency RBCs, (B) sphero-echinocytes, and (C) spherocytes over the entire curved channel (0° < *θ* < 180°) in each corresponding binary suspensions of normal RBCs with aberrant RBCs.

These results demonstrate that in the curved tube, not only do we see margination of aberrant cells as found in the straight tube, but very strong localization of the marginated cells on the centerplane. The mechanism of this localization originates in the *θ*-dependence of the cell-free layer thickness, as illustrated in Fig. 5. Fig.5(A) shows a simulation snapshot of cross-section cell distribution at *ϕ* = *π*/2, in which aberrant cells are highly localized near the outer side centerplane. We noted above that the CFL thickness is approximately uniform for *π*/2 *< θ <* 3*π*/2 – i.e. along the inner wall, but on the outer side, the CFL thickness increases, reaching a maximum on the centerplane on the outer wall – i.e. the CFL thickness increases as *θ* → 0. A marginated cell on the outer wall will experience more collisions from the side with the thinner CFL than the thicker, thus being driven on average toward the region where the CFL is thickest, *θ* = 0. Fig.5(B) shows the trajectories of marginated aberrant RBCs on the *θ* − *ϕ* plane, demonstrating that as *ϕ* increases, *θ* tends to decrease and aberrant cells move towards the centerplane. We illustrate this mechanism schematically in Fig.5(C). Finally, we must address why there is localization along the centerplane on the inner wall (*θ* = *π*). This results from the simple fact that the outer wall over half a wavelength of the curved shape is the inner wall over the other half; the CFL thickness is nearly constant along the inner wall, driving no net motion in *θ*, and cells driven toward *θ* = 0 on the outer wall tend to remain there while moving along the inner wall.

**Fig. 5.**
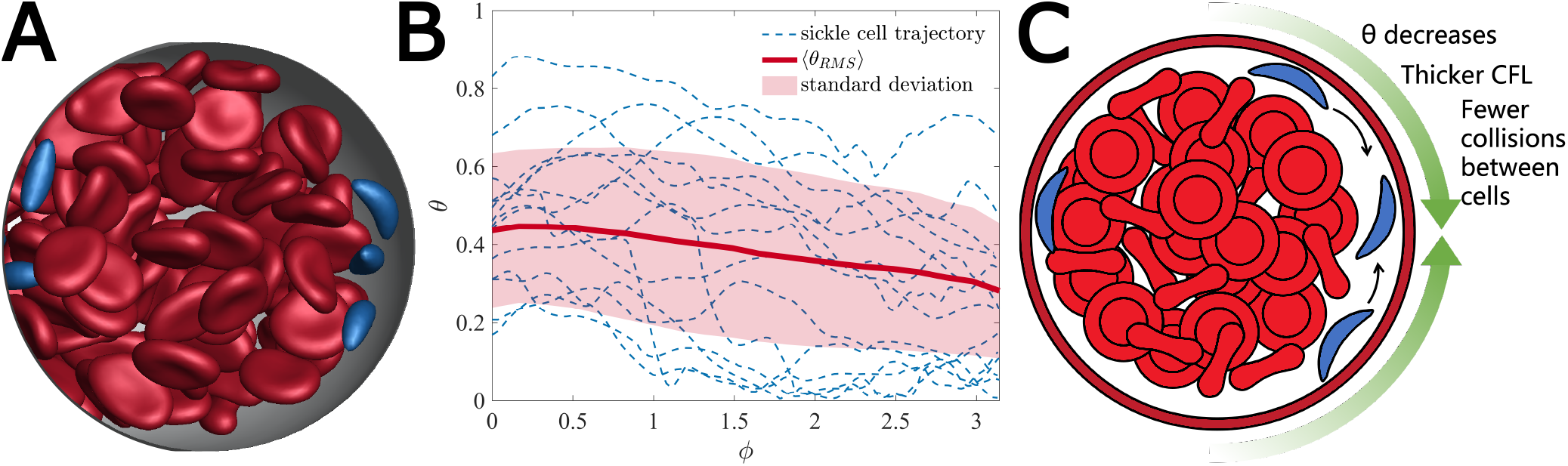
(A) Simulation snapshot showing the cross-section cell distribution (*ϕ* = *π*/2). (B) Trajectories of sickle RBCs on the *θ* − *ϕ* plane. Dashed blue curves are individual trajectories; the red line denotes the root mean squared trajectory of the blue curves, and the red shaded area is for the corresponding standard deviation. (C) A schematic mechanism for localization of marginated cells to *θ* = 0.

Fig.6(*A*) shows snapshots of the spatial distribution of excess wall shear stress 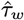 in SCD suspensions; results for the other blood disorders can be found in the SI. The presence of a sickle RBC close to the wall directly causes local fluctuations in wall shear stress, as can be observed from the transparent view in Fig.6(*B*).

To capture the spatial dependence of RBC-induced wall shear stress 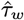,, the probability density profiles of *τ*_*w*_ in SCD and healthy RBC suspensions over the different *θ* − areas on the vascular surface are presented in Fig.6(C). In the healthy RBC suspension, the probability of high 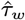 is largest near the inner side (3*π*/4 < *θ* < *π*), followed by the intermediate area of (*π*/4 < *θ* < 3*π*/4), and smallest over the outer side (0 < *θ* < *π*/4), consistent the fact that the CFL is thinnest near the inner side and thickest near the outer side. As in the straight tube, it is observed that the cases with aberrant RBCs exhibit a distinct excess of large positive fluctuations, again attributable to the margination of these small stiff cells to the vessel wall. Furthermore, the disparity at high RBC-induced wall shear stress 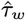 of the probability distribution profiles between diseased and healthy RBC suspensions is most pronounced at the outer side, which implies that the localization of the marginated aberrant RBCs to the center plane elevates the probability of high additional wall shear stress by an order of magnitude over the outer side wall of the curved tube.

**Fig. 6.**
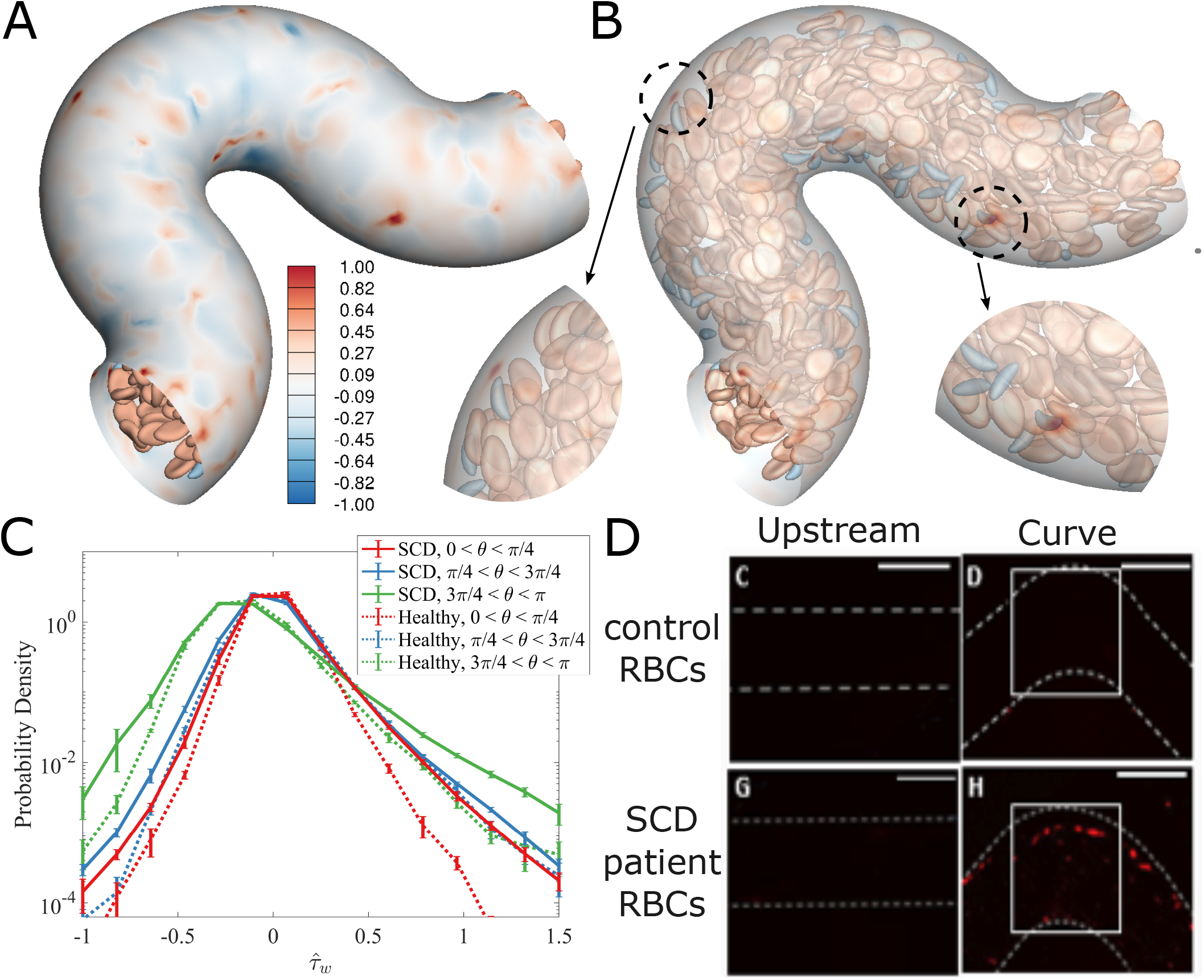
(A) Simulation snapshots and (B) corresponding transparent views of additional wall shear stress 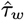 induced by the presence of the cells in suspensions of normal RBCs with sickle RBCs within the curved channel. The color on the vascular surface denotes the RBC-induced wall shear stress strength 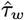. (C) The time-averaged probability distribution of the additional wall shear stress 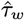 for SCD RBC suspension and healthy RBC suspension over the different *θ* − area on the curved vascular surface. (D) Mechanically stiff SCD RBCs upregulate E-selectin at the curvature site of vasculature models, indicating pro-inflammatory endothelial signaling (10). Scale bars = 200*μm*

The geometries of blood vessels have been found to play a role in the development of endothelial dysfunction in blood disorders. Using an endothelialized microfluidic model, Wang et al.(10) discovered that endothelial cells exposed to SCD RBCs exhibited an increase in endothelial inflammation along the outside wall of the bend in the curved regions of vessels. This result is illustrated in Fig.6(D), which shows expression levels of E-selectin, which is upregulated when cells are in a pro-inflammatory state, upstream and in a curved segment of endothelialized microfluidic channels through which RBC suspensions have flowed. While the channel sizes in the experiments are much larger than those simulated here, the qualitative pattern of endothelial inflammation there is consistent with the margination patterns we observe here.

## Conclusions

Blood disorders lead to changes in red blood cell size, shape, and stiffness, and thus to changes in how aberrant cells are distributed in the cross-section of blood vessels, and changes in the interaction between cells and blood vessels. Inflammation and dysfunction of endothelial cells lining blood vessels are associated with the risk of pathophysiologic complications like stroke and atherosclerosis.

This study describes results from detailed cell-level simulations of blood in straight and serpentine tubes, addressing the hypothesis that the margination of aberrant cells leads to substantial changes in the local shear stress environment of the blood vessel wall, possibly contributing to the observed dysfunction and inflammation. We compare cell distributions and wall shear stress profiles between suspensions of normal blood and blood containing aberrant RBCs that model sickle cell disease (SCD), iron deficiency anemia, COVID-19, and spherocytosis. In all cases, the smaller and stiffer aberrant RBCs marginate and increase the fluctuations in wall shear stress. Probability density profiles of wall shear stress show that cases with aberrant RBCs display a significantly higher probability of high wall shear stress than in suspensions of healthy cells. The difference is most notable in COVID-19 and SCD RBC suspensions.

In the serpentine curved tube case, the marginated aberrant cells tend to marginate most strongly to the symmetry plane of the channel and the outer side of the curved tube, becoming very strongly localized in those regions. This result implies the possibility of strongly localized endothelial damage in blood vessels with complex geometries. These findings highlight the importance of considering vascular geometry and the presence of aberrant RBCs in the development of vasculopathy.

Overall, our study indicates that the biophysical alterations of red cells in various disorders, in and of themselves, can directly alter the shear stress the underlying endothelium is exposed to. This suggests that that red cell biophysics and the pathologic changes thereof may directly affect endothelial mechanobiological pathways, which, in turn, may be associated with chronic endothelial inflammation or dysfunction that lead to disorders such cardiovascular disease and stroke.

Experiments to complement these computational results are now being conducted in our group. In addition, our study also suggests that clinically, more attention should be paid to the biophysical alterations of the red cells themselves, which currently are viewed as hallmarks of the associated disease but not necessarily as biomarkers per se; our work suggests otherwise and indicates that the red cell shape, size, deformability should be re-examined more rigorously as potential correlates or predictors of clinical endpoints. Finally, our work indicates that therapeutic interventions that “fix” or remove the biophysical alterations of red cells should be explored for the associated diseases. Indeed red cell pheresis for various hematology conditions may improve the vascular dysfunction associated with those diseases and for sickle cell disease in particular, recently FDA-approved treatments that improve “red cell health” may also alleviate vasculopathy.

## Materials and Methods

### Model set-up

We consider a flowing suspension of RBCs, which we model as deformable fluid-filled elastic capsules, in rigid straight (Fig.1) and curved “serpentine” (Fig.3) cylindrical tubes with radius *R* = 20*μ*m. No-slip boundary conditions are imposed on the walls of the tube, while periodic boundary conditions are applied in the flow direction. The suspension is subjected to a unidirectional pressure-driven flow, and the velocity field in the absence of RBCs field within the straight cylindrical tube is given by the Poiseuille flow. In this study, the flow is driven by a constant pressure gradient, which is equivalent to fixing the mean wall shear rate at 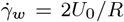 where *U*_0_ is the undisturbed centerline velocity. For the curved cylindrical channel, the pressure gradients are determined using defining an equivalent straight cylindrical channel with the same centerline length and radius.

This study considers both homogeneous and heterogeneous suspensions of various components, including normal RBCs and aberrant RBCs (e.g., iron deficiency RBCs, sickle RBCs, spheroechinocytes, and spherocytes). In binary suspensions, normal RBCs are generally considered primary components (denoted as “p”), while aberrant RBCs as trace components (denoted as “t”). A normal RBC is modeled as a flexible capsule having the spontaneous shape being a biconcave discoidal for shear elasticity and an oblate spheroid for bending elasticity (41, 42), with a radius of *a* = 4*μm*. The idRBCs have the same rest shape as normal RBCs, except the radius of idRBCs is 0.76*a* (12). The rest shapes of sickle RBCs, spherocytes, and sphero-echinocytes are curved oblate, spherical with a diameter of 5*μm*, and spiked spherical, respectively. The cell membranes are modeled as an isotropic and hyperelastic surface with interfacial shear modulus *G*, incorporating shear elasticity, area dilatation, volume conservation, and bending resistance. Details of the membrane mechanics model and validation against experimental observations are given in (41).

The deformability of a capsule in pressure-driven flow is measured by the dimensionless capillary number 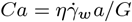. In this study, *G* of the aberrant RBCs is assumed to be five times that of normal RBCs, which leads to that *Ca*_*t*_ for aberrant RBCs is always around less 0.2 times that of *Ca*_*p*_ for biconcave discoid RBCs. In this study, Ca_*p*_ is set to 1.0 for normal RBCs, Ca_*t*_ is 0.15 for idRBCs, 0.20 for sickle RBCs, 0.15 for sphero-echinocytes, and 0.125 for spherocytes, which corresponds to 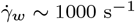. In the binary suspension, the number fractions for normal RBCs *X*_*b*_ is set to 0.9, and for aberrant RBCs, *X*_*t*_ is 0.1, so the overall number density ratio *n*_*p*_/*n*_*t*_ = 9. In this study, the total cell volume fraction (hematocrit) is set to be *ϕ* ≈ 0.20. To simplify the computations in this initial study, the suspending fluid and the fluid inside the cells are assumed to have the same viscosity.

In our simulation, the particle Reynolds number, defined as 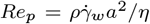, is set to be 0.1, and the fluid is assumed to be incompressible and Newtonian; therefore the flow is governed by the Navier-Stokes and continuity equations. A projection method is used to advance the velocity field in time. The straight tube is embedded in a cuboidal computational domain with the size of 10*a* × 10*a* × 10*a*, and an Eulerian grid of 100 × 100 × 100 is used. For the serpentine channel case, the cuboidal computational domain’s size is 32*a* × 26*a* × 10*a*, and the Eulerian grid of 320 × 260 × 100 is used. The immersed boundary method (IBM) is used to handle fluid-structure interaction. Specifically, the current model considers two types of immersed boundaries: deformable moving cellular membranes and rigid nonmoving vascular walls. The capsule membrane is discretized into *N*_Δ_ piecewise flat triangular elements; *N*_Δ*p*_ = 1280 for normal RBC, while *N*_Δ*t*_ = 682 for sickle RBC, *N*_Δ*t*_ = 816 for idRBC, *N*_Δ*t*_ = 620 for spherocyte, and *N*_Δ*t*_ = 1134 for sphero-echinocyte. Different *N*_Δ_ are chosen to ensure that the triangular elements on both capsules are close in size. We use “continuous forcing” IBM and “direct forcing” IBM methods for the RBC membranes and tube wall, respectively. The numerical methodology follows the approach described in (43, 44).

## ACKNOWLEDGMENTS

This work was supported by National Science Foundation grant CBET-2042221 (M.D.G., X.C.), an American Society of Hematology (ASH) Research Training Award for Fellows (RTAF), National Institutes of Health, Institute of Heart, Lung, and Blood (NIH NHLBI) grant T32HL139443 and Pediatric Loan Repayment Program (LRP) Award L40HL149069 (C.C.), National Institutes of Health, National Heart, Lung, and Blood Institute grant R35HL145000 (W.A.L.). The work was performed in part at the Georgia Tech Institute for Electronics and Nanotechnology, a member of the National Nanotechnology Coordinated Infrastructure (NNCI), which is supported by the National Science Foundation grant ECCS-2025462. This work used the Advanced Cyberinfrastructure Coordination Ecosystem: Services & Support (ACCESS). In particular, it used the Expanse system at the San Diego Super-computing Center (SDSC) through allocation MCB190100.

## References

1. C Hahn, MA Schwartz, Mechanotransduction in vascular physiology and atherogenesis. Nat. reviews Mol. cell biology 10, 53–62 (2009).

2. BK Walther, et al., Mechanotransduction-on-chip: vessel-chip model of endothelial yap mechanobiology reveals matrix stiffness impedes shear response. Lab on a Chip 21, 1738–1751 (2021).

3. KC Wang, et al., Flow-dependent yap/taz activities regulate endothelial phenotypes and atherosclerosis. Proc. Natl. Acad. Sci. 113, 11525–11530 (2016).

4. M He, M Martin, T Marin, Z Chen, B Gongol, Endothelial mechanobiology. APL bioengineering 4, 010904 (2020).

5. T Panciera, L Azzolin, M Cordenonsi, S Piccolo, Mechanobiology of yap and taz in physiology and disease. Nat. reviews Mol. cell biology 18, 758–770 (2017).

6. X Zhou, W Xu, Y Xu, Z Qian, Iron supplementation improves cardiovascular outcomes in patients with heart failure. The Am. journal medicine 132, 955–963 (2019).

7. C Souilhol, et al., Endothelial responses to shear stress in atherosclerosis: a novel role for developmental genes. Nat. Rev. Cardiol. 17, 52–63 (2020).

8. E Tzima, et al., A mechanosensory complex that mediates the endothelial cell response to fluid shear stress. Nature 437, 426–431 (2005).

9. R Mannino, et al., Vascular geometry and flow profile mediate pathological cell-cell interactions in sickle cell disease as measured with” do-it-yourself”“ endothelial-ized” microfluidics (2014).

10. Y Wang, et al., Vessel geometry interacts with red blood cell stiffness to promote endothelial dysfunction in sickle cell disease. Blood 126, 965 (2015).

11. JF Bertles, PF Milner, et al., Irreversibly sickled erythrocytes: a consequence of the heterogeneous distribution of hemoglobin types in sickle-cell anemia. The J. Clin. Investig. 47, 1731–1741 (1968).

12. C Caruso, et al., Pathologic mechanobiological interactions between red blood cells and endothelial cells directly induce vasculopathy in iron deficiency anemia. IScience 25, 104606 (2022).

13. SM Recktenwald, et al., Cross-talk between red blood cells and plasma influences blood flow and omics phenotypes in severe covid-19. Elife 11, e81316 (2022).

14. S Druzak, et al., Multiplatform analyses reveal distinct drivers of systemic pathogenesis in adult versus pediatric severe acute COVID-19. Nat. Commun. 14, 1638 (2023).

15. S Perrotta, PG Gallagher, N Mohandas, Hereditary spherocytosis. The Lancet 372, 1411– 1426 (2008).

16. JC Firrell, HH Lipowsky, Leukocyte margination and deformation in mesenteric venules of rat.

17. GJ Tangelder, HC Teirlinck, DW Slaaf, RS Reneman, Distribution of blood platelets flowing in arterioles. Am. J. Physiol. Circ. Physiol. 248, H318–H323 (1985).

18. JJ Bishop, AS Popel, M Intaglietta, PC Johnson, Effect of aggregation and shear rate on the dispersion of red blood cells flowing in venules. Am. J. Physiol. Circ. Physiol. 283, H1985– H1996 (2002).

19. A Kumar, MD Graham, Segregation by membrane rigidity in flowing binary suspensions of elastic capsules. Phys. Rev. E 84, 066316 (2011).

20. A Kumar, RGH Rivera, MD Graham, Flow-induced segregation in confined multicomponent suspensions: effects of particle size and rigidity. J. Fluid Mech. 738, 423–462 (2014).

21. K Sinha, MD Graham, Shape-mediated margination and demargination in flowing multicomponent suspensions of deformable capsules. Soft matter 12, 1683–1700 (2016).

22. QM Qi, ESG Shaqfeh, Theory to predict particle migration and margination in the pressure-driven channel flow of blood. Phys. Rev. Fluids 2, 093102 (2017).

23. K Mueller, DA Fedosov, G Gompper, Understanding particle margination in blood flow - A step toward optimized drug delivery systems. Med. Eng. & Phys. 38, 2 10 (2016-01).

24. A Kumar, MD Graham, Mechanism of Margination in Confined Flows of Blood and Other Multicomponent Suspensions. Phys. Rev. Lett. 109, 108102 (2012).

25. V Narsimhan, H Zhao, ESG Shaqfeh, Coarse-grained theory to predict the concentration distribution of red blood cells in wall-bounded Couette flow at zero Reynolds number. Phys. Fluids 25, 061901 (2013).

26. RGH Rivera, K Sinha, MD Graham, Margination Regimes and Drainage Transition in Confined Multicomponent Suspensions. Phys. Rev. Lett. 114, 188101 (2015).

27. RGH Rivera, X Zhang, MD Graham, Mechanistic theory of margination and flow-induced segregation in confined multicomponent suspensions: Simple shear and Poiseuille flows*. Phys. Rev. Fluids 1, 060501 (2016).

28. C Collins, et al., Haemodynamic and extracellular matrix cues regulate the mechanical phenotype and stiffness of aortic endothelial cells. Nat. communications 5, 3984 (2014).

29. J Eyckmans, T Boudou, X Yu, CS Chen, A hitchhiker’s guide to mechanobiology. Dev. Cell 21, 35–47 (2011).

30. D Harrison, et al., Endothelial mechanotransduction, nitric oxide and vascular inflammation. J. internal medicine 259, 351–363 (2006).

31. DE Ingber, From mechanobiology to developmentally inspired engineering. Philos. Transactions Royal Soc. B: Biol. Sci. 373, 20170323 (2018).

32. D Ingber, Mechanobiology and diseases of mechanotransduction. Annals medicine 35, 564– 577 (2003).

33. Ji Abe, BC Berk, Novel mechanisms of endothelial mechanotransduction. Arter. thrombosis, vascular biology 34, 2378–2386 (2014).

34. C Caruso, et al., Stiff erythrocyte subpopulations biomechanically induce endothelial inflammation in sickle cell disease. Blood 134, 3560 (2019).

35. X Zhang, C Caruso, WA Lam, MD Graham, Flow-induced segregation and dynamics of red blood cells in sickle cell disease. Phys. Rev. Fluids 5, 053101 (2020).

36. H Flyvbjerg, HG Petersen, Error estimates on averages of correlated data. The J. Chem. Phys. 91, 461–466 (1989).

37. B Klitzman, BR Duling, Microvascular hematocrit and red cell flow in resting and contracting striated muscle. Am. J. Physiol. Circ. Physiol. 237, H481–H490 (1979).

38. IH Sarelius, BR Duling, Direct measurement of microvessel hematocrit, red cell flux, velocity, and transit time. Am. J. Physiol. Circ. Physiol. 243, H1018–H1026 (1982).

39. J Dupire, M Socol, A Viallat, Full dynamics of a red blood cell in shear flow. Proc. Natl. Acad. Sci. 109, 20808–20813 (2012).

40. TM Fischer, Shape memory of human red blood cells. Biophys. journal 86, 3304–3313 (2004).

41. K Sinha, MD Graham, Dynamics of a single red blood cell in simple shear flow. Phys. Rev. E 92, 042710 (2015).

42. E Evans, YC Fung, Improved measurements of the erythrocyte geometry. Microvasc. research 4, 335–347 (1972).

43. P Balogh, P Bagchi, A computational approach to modeling cellular-scale blood flow in complex geometry. J. Comput. Phys. 334, 280–307 (2017).

44. R Mittal, et al., A versatile sharp interface immersed boundary method for incompressible flows with complex boundaries. J. computational physics 227, 4825–4852 (2008).

