## Supplementary material for "Marginated aberrant red blood cells induce pathologic vascular stress fluctuations in a computational model of hematologic disorders": SI Appendix

#### This PDF file includes:

Supporting text
Figs. S1 to S12
Legends for Movies S1 to S8
SI References

#### Other supporting materials for this manuscript include the following:

Movies S1 to S8

### Supporting Information Text

### Methods

We have developed a computational model utilizing the immersed boundary method (IBM) to investigate cellular blood flow in complex vessel geometries. This approach offers the advantage of modeling flows in arbitrary geometries and accurately resolving the large deformation and dynamics of blood cells. Our model considers blood as a confined flowing suspension of red blood cells (RBCs) in the plasma. Two distinct types of boundaries are involved: deformable cellular membranes and rigid non-moving vascular walls with complex geometry. To handle these two types of interfaces, we use the continuous-forcing and direct-forcing immersed boundary methods, respectively. Specifically, the continuous-forcing IBM method couples surface stresses on the RBC membrane with the fluid flow, while the sharp-interface ghost-node immersed boundary method (GNIBM) is used to treat the rigid, non-moving vascular walls (1, 2). The flow solver is based on a coupled finite-volume/spectral method, and our IBM code is parallelized using a hybrid MPI/OpenMP strategy.

**Governing Equation** Assuming that blood plasma is both Newtonian and incompressible, the governing equations for its flow can be expressed as the incompressible Navier-Stokes equations.

$$
\begin{gathered}
Re\left(\frac{\partial \boldsymbol{u}}{\partial t}+\nabla\cdot\boldsymbol{u}\boldsymbol{u}\right)=-\nabla P+\nabla^2\boldsymbol{u}+\boldsymbol{F}\\
\nabla\cdot\boldsymbol{u}=0
\end{gathered}
\tag{1}
$$

**Flow Solver** The Chorin projection method is utilized to advance the velocity field $\boldsymbol{u}$. This method involves solving an advection-diffusion equation (ADE) to determine the intermediate velocity field $\boldsymbol{u}^*$, followed by solving a Poisson equation for pressure $P$ to enforce the divergence-free constraints.

$$
\begin{gathered}
Re\frac{\boldsymbol{u}^*-\boldsymbol{u}^n}{\Delta t}=\nabla^2\boldsymbol{u}-Re\nabla\cdot\boldsymbol{u}\boldsymbol{u}+\boldsymbol{F}\\
\nabla^2 P=\frac{Re}{\Delta t}\nabla\cdot\boldsymbol{u}^*\\
\frac{\boldsymbol{u}^{n+1}-\boldsymbol{u}^*}{\Delta t}=-\frac{1}{Re}\nabla P
\end{gathered}
\tag{2}
$$

For the ADE, we treat both the convection term and the body force term with the 2nd-order Adams-Bashforth method (AB2), and the diffusion term with the Crank-Nicholson method for numerical stability.

$$
Re\frac{\boldsymbol{u}^*-\boldsymbol{u}^n}{\Delta t}=0.5\nabla^2(\boldsymbol{u}^*+\boldsymbol{u}^n)+1.5N\left(\boldsymbol{u}^n\right)-0.5N\left(\boldsymbol{u}^{n-1}\right)+1.5\boldsymbol{F}^n-0.5\boldsymbol{F}^{n-1}
\tag{3}
$$

where $N(\boldsymbol{u})=-Re\nabla\cdot\boldsymbol{u}\boldsymbol{u}$ denotes the nonlinear convection, evaluated then by the central differencing. To leverage the efficient inversion of tri-diagonal matrices, a Locally One-Dimensional (LOD) Alternating Direction Implicit (ADI) scheme is employed to solve the ADE. This scheme involves four steps: first, the explicit terms are handled; then, the x, y, and z directions are solved implicitly, one at a time.

$$
\begin{gathered}
\frac{Re}{\Delta t}\boldsymbol{u}^{*/4}=\frac{Re}{\Delta t}\boldsymbol{u}^n+1.5N\left(\boldsymbol{u}^n\right)-0.5N\left(\boldsymbol{u}^{n-1}\right)+1.5\boldsymbol{F}^n-0.5\boldsymbol{F}^{n-1}\\
\left(\frac{Re}{\Delta t}-\frac{1}{2}\frac{\partial^2}{\partial x^2}\right)\boldsymbol{u}^{**/4}=\left(\frac{Re}{\Delta t}+\frac{1}{2}\frac{\partial^2}{\partial x^2}\right)\boldsymbol{u}^{*/4}\\
\left(\frac{Re}{\Delta t}-\frac{1}{2}\frac{\partial^2}{\partial y^2}\right)\boldsymbol{u}^{***/4}=\left(\frac{Re}{\Delta t}+\frac{1}{2}\frac{\partial^2}{\partial y^2}\right)\boldsymbol{u}^{**/4}\\
\left(\frac{Re}{\Delta t}-\frac{1}{2}\frac{\partial^2}{\partial z^2}\right)\boldsymbol{u}^{*}=\left(\frac{Re}{\Delta t}+\frac{1}{2}\frac{\partial^2}{\partial z^2}\right)\boldsymbol{u}^{***/4}
\end{gathered}
\tag{4}
$$

To solve the 3D pressure Poisson equation (PPE), we assume that the computational domain is periodic in two directions, namely $x$ and $y$. We begin by performing a 2D fast Fourier transform (FFT) on each $x-y$ plane. Next, solve for the Fourier coefficients along the z direction. Finally, perform a 2D inverse fast Fourier transform (iFFT) on each $x-y$ plane to obtain the pressure field.

**Spatial Discretization** The IBM requires an Eulerian representation for the fluid flow and a Lagrangian representation for the immersed boundaries such as the RBC membrane. The governing equations are solved on the Eulerian grid. To avoid odd-even decoupling, spatial discretization is based on the staggered mesh, in which these scalar variables, such as pressure $P$, are located at the grid center, while vector variables, such as velocity $u$ and force $F$, are located at the grid faces. All spatial derivatives are evaluated using second-order differencing. As for the Lagrangian mesh on the immersed boundaries, the membrane is discretized into piecewise flat triangular elements. An open-source software Gmsh (3) is used to generate the surface mesh for both vessels (Fig. S1) and various red blood cells (Fig. S2).

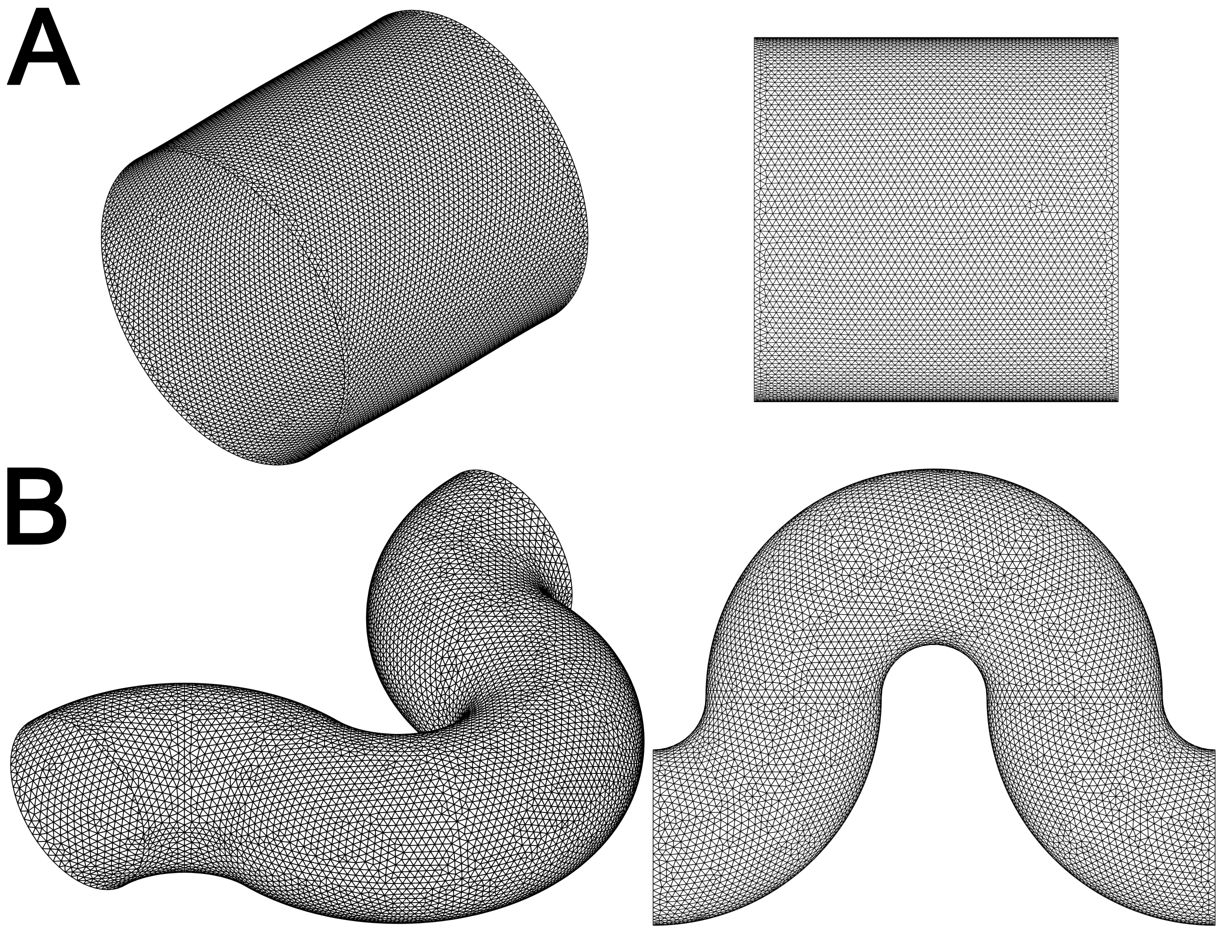


**Fig. S1.** Surface mesh for (A) a straight cylindrical tube (radius $R = 20\mu m$ and length $L = 40\mu m$) and (B) a curved (serpentine) cylindrical channel (major radius $R_1 = 32\mu m$ and minor radius $R_2 = 20\mu m$).

**Membrane Mechanics** The RBC membrane is assumed to resist shear deformation, area dilatation, volume conservation(4), and bending resistance. The total strain energy of the RBC membrane $S$ is given by,

$$E = \frac{K_B}{2} \int_S (2\kappa_H + c_0)^2 \, dS + \overline{K_B} \int_S \kappa_G dS + \int_S W dS$$

where $K_B$ and $\overline{K_B}$ are the bending moduli, and $W$ is the shear strain energy density; $\kappa_H, \kappa_G$ are the mean and Gaussian curvature of the surface, respectively; $c_0 = -2H_0$ is the spontaneous curvature, where $H_0$ is the mean curvature of the spontaneous shape. The first two terms correspond to the Canham-Helfrich bending energy (5, 6), and the third term comes from the shear strain energy stored in the RBC membrane. The strain energy density is computed based on the Skalak model (7). A finite element method (FEM) is developed to find the surface stress as a result of membrane deformation (8).

**Continuous Forcing IBM** The cellular membrane is deformed by the fluid flow, while the flow is altered by the membrane deformation in turn. This fluid-structure interaction (FSI) between flow and membrane is characterized via the continuous forcing IBM. The idea of this method is to add an external force term $\boldsymbol{F}$ to the right-hand side of the Navier-Stokes equation. The external force term originates from the membrane stress $f_{\text{membrane}}$. A discretized delta function is used to spread the singular force on the membrane to the surrounding fluid and interpolate the fluid velocity back to the membrane.

$$\begin{aligned} \boldsymbol{F} &= \int_S \boldsymbol{f}_{\text{membrane}} \, \delta\left(\boldsymbol{x} - \boldsymbol{x}'\right) d\boldsymbol{x}' \\ \boldsymbol{u}_{\text{membrane}} &= \int_{\mathcal{T}} \boldsymbol{u} \delta\left(\boldsymbol{x}' - \boldsymbol{x}\right) d\boldsymbol{x} \end{aligned} \tag{5}$$

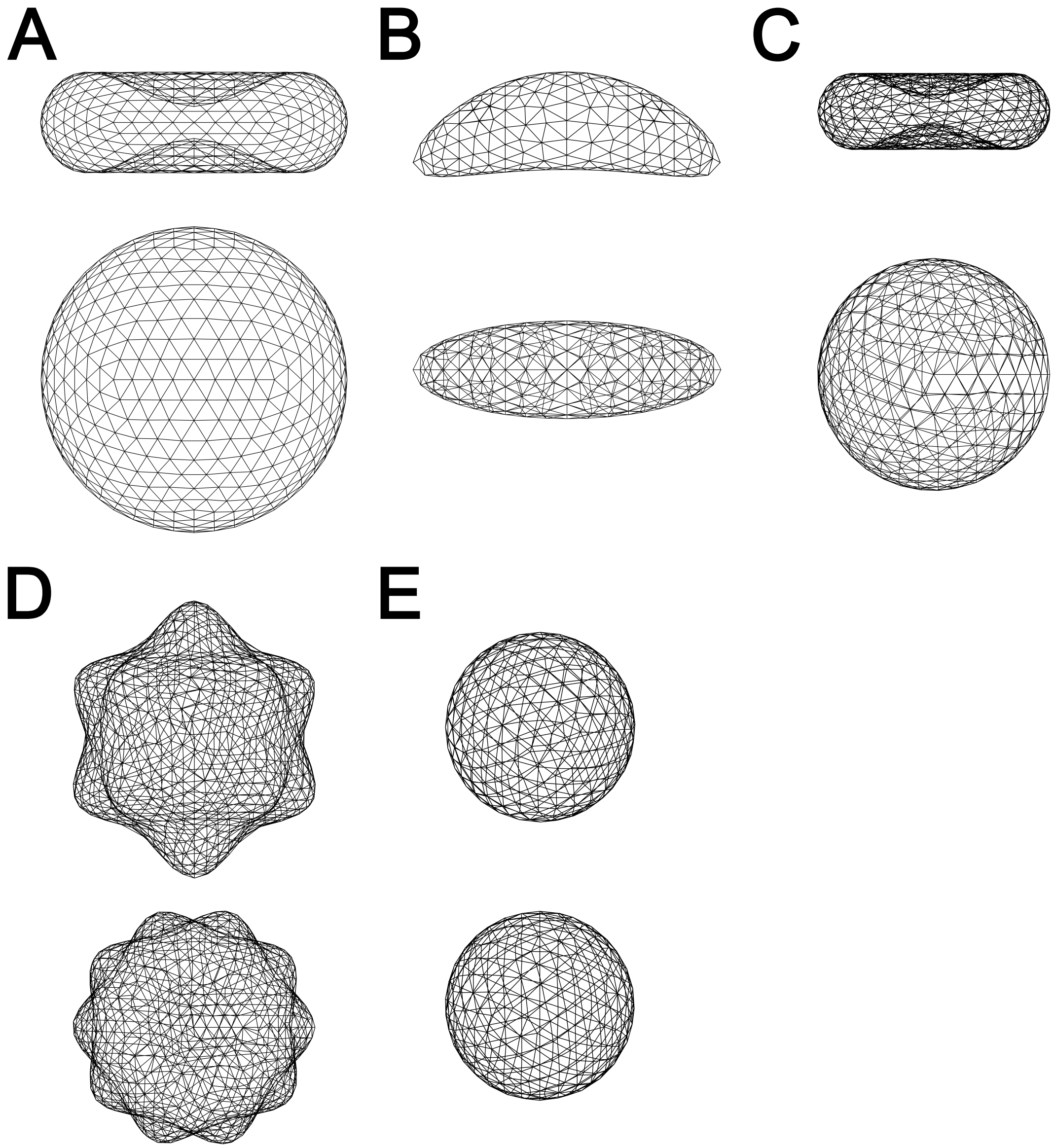


**Fig. S2.** The surface mesh for capsules of (A) biconcave-discoid normal RBC, (B) sickle RBC, (C) iron deficiency RBC, (D) sphere-echinocyte, and (E) spherocyte. Note the size of each cell species in this figure is corresponding to the cell used in our simulation. The radius of normal RBC is $4\mu m$; aberrant RBCs are smaller than normal RBCs.

$\delta$ is the three-dimensional Dirac-delta function, and $\mathbf{x}$ and $\mathbf{x}'$ are the locations in the flow domain $\mathcal{T}$ and on the cell surface $S$, respectively. A numerical approximation of the delta function is chosen to be

$$\delta\left(\mathbf{x}-\mathbf{x}'\right)=\begin{cases}\dfrac{1}{64\Delta^3}\displaystyle\prod_{i=1}^{3}\left[1+\cos\frac{\pi}{2\Delta}\left(x_i-x_i'\right)\right], & |x_i-x_i'|\le 2\Delta\\ 0, & |x_i-x_i'|>2\Delta\end{cases} \quad [6]$$

where $\Delta$ is the Eulerian grid size.

**Direct Forcing IBM** Direct forcing IBM is used to treat another type of immersed boundary, the rigid, non-moving but geometrically complex vessel walls. Specifically, GNIBM is used to impose no-slip velocity boundary conditions on the vessel surface (1). The idea of this method is to modify these differential operators appropriately to satisfy the no-slip boundaries condition on the geometrically-complex vascular surface while maintaining second-order accuracy.

**Indicator Function for Hematocrit Analysis** To characterize the cellular spatial distribution in RBC suspension, we define an indicator function $I(\mathbf{x},t)$, such that the indicator function is one inside a cell, and zero outside a cell. It can be shown (9) that the indicator function $I(\mathbf{x},t)$ follows a Poisson equation as

$$\nabla^2 I=\nabla\cdot\mathbf{G},\quad \mathbf{G}(\mathbf{x},t)=\int_S\delta\left(\mathbf{x}-\mathbf{x}'\right)\mathbf{n}dS \quad [7]$$

where the $\mathbf{G}(\mathbf{x},t)$ is an Eulerian variable constructed from the cell surface normals $\mathbf{n}$.

**Wall Shear Stress Evaluation** To compute wall shear stress, the traction vector $\mathbf{t} = \boldsymbol{\tau}\cdot\mathbf{n}$ at the wall is determined using the velocity field approach outlined in (10, 11), where $\boldsymbol{\tau}$ is the stress tensor and $\boldsymbol{n}$ is the unit normal vector. The no-slip condition on the blood vessel surface yields the expression for the local wall shear stress $t_s = \mu\partial u_s/\partial r$. Note $s$ denotes the local stream-wise direction. The second-order differencing method is utilized to numerically evaluate velocity derivatives.

### Model Validation

**Elastic Spherical Capsule in Simple Shear Flow** We consider a moving deformable spherical capsule subjected to simple shear flow, see Fig.S3(A). The deformation of a capsule will cause stretching stress and bending stress on its surface. The deformability of a capsule is characterized by the nondimensional Capillary number $\mathrm{Ca} = \mu\dot{\gamma}a/G$ and the nondimensional bending modulus $\hat{\kappa}_B = K_B/a^2G$, where $\mu$ is the fluid viscosity, $\dot{\gamma}$ is shear rate, $a$ is the radius of the capsule, $G$ is the shear elasticity modulus and $K_B$ is the bending modulus. Larger $Ca$ and $\hat{\kappa}_B$ means that a capsule is more flexible, while a stiffer capsule has smaller $Ca$ and $\hat{\kappa}_B$. The deformation of the capsule is described by the Taylor shape parameter defined as $D_{xz} = (L-B)/(L+B)$, where $L$ and $B$ are the maximum and minimum radial distances of an ellipsoid with the same inertia tensor. In Fig.S3(B), we show that steady state values of Taylor deformation parameter $D$ as a function of dimensionless Capillary number Ca for different $\hat{\kappa}_B$. Good agreement is found between our numerical results and simulation results from previous literature (12).

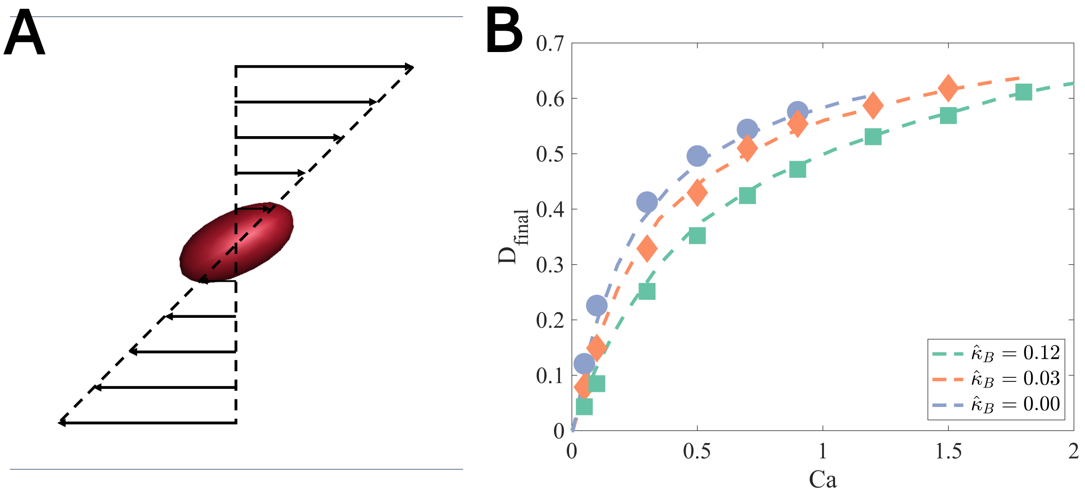


**Fig. S3.** (A) Schematic diagram of an elastic spherical capsule in simple shear flow. Velocity Dirichlet boundary conditions are imposed on both the top and bottom plates to create a simple shear flow. Because of the existence of a constant shear rate, the sphere capsule is stretched and then deformed into an ellipsoid. (B) Steady-state Taylor deformation parameter for a spherical capsule as a function of $Ca$. Dash lines are results from previous literature (12), and symbols are our simulation results.

**Stationary Rigid Sphere in Simple Shear Flow** We consider a rigid stationary sphere subjected to a linear shear flow, which is given by $\mathbf{u}^\infty = [\dot{\gamma}z, 0, 0]$. The schematic of the model set is shown in Fig.S4(A). The analytical solution for this problem is given by

$$\begin{aligned} u &= \frac{y\dot{\gamma}}{2}\left[1-\left(\frac{r}{a}\right)^{-5}\right] + \frac{y\dot{\gamma}}{2}\left[1-\left(\frac{r}{a}\right)^{-3}\right] - \frac{5}{2}\left(\frac{x}{a}\right)^{2} y\dot{\gamma}\left[\left(\frac{r}{a}\right)^{-5} - \left(\frac{r}{a}\right)^{-7}\right] \\ v &= \frac{x\dot{\gamma}}{2}\left[1-\left(\frac{r}{a}\right)^{-5}\right] - \frac{x\dot{\gamma}}{2}\left[1-\left(\frac{r}{a}\right)^{-3}\right] - \frac{5}{2}\left(\frac{y}{a}\right)^{2} x\dot{\gamma}\left[\left(\frac{r}{a}\right)^{-5} - \left(\frac{r}{a}\right)^{-7}\right] \\ w &= -\frac{5}{2}\frac{xyz\dot{\gamma}}{a^2}\left[\left(\frac{r}{a}\right)^{-5} - \left(\frac{r}{a}\right)^{-7}\right] \end{aligned} \qquad [8]$$

where $u, v, w$ are the velocity component in each dimension, $r = \sqrt{x^2+y^2+z^2}$ is the distance from a point in flow to the center of the sphere, $a$ is the radius of the sphere. Using the direct forcing IBM, we set the velocity on the sphere surface to zero (no-slip boundary condition). Note the solution above is for unbounded shear flow, thus we impose the Dirichlet velocity boundary conditions on both the upper and bottom walls and choose a larger computation domain to reduce the error by periodic boundaries. After the velocity field reaches its steady state, our numerical results are then compared with the analytical solutions. $L_1$ and $L_2$ error norms for different velocity components $u, v, w$ are plotted in Fig. S4B. It is found that the direct forcing IBM in our model presents second-order accuracy.

### Impact of Membrane Curvature on RBC Dynamics

Although efforts to understand RBC dynamics numerically have spanned the past two decades, the majority of these works have focused on RBC dynamics in cases where the RBC shape is symmetric across the shear plane or where the dimple is centered

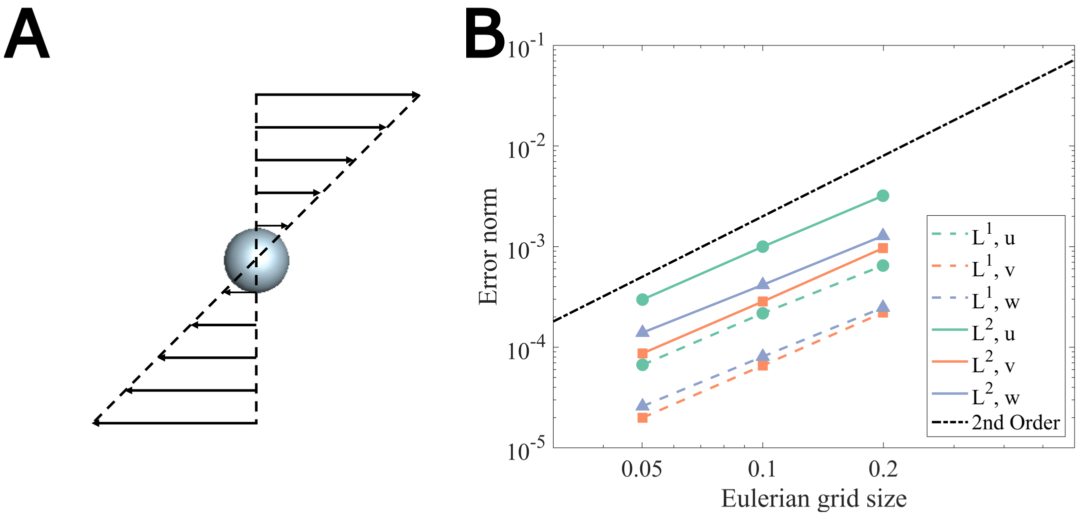


**Fig. S4.** (A) Schematic of simulating a rigid stationary sphere subjected to a linear shear flow. The velocity field on the top and bottom wall is set to be the analytical solution (Dirichlet boundary condition). Periodic boundary conditions are imposed along $x$ and $y$ direction. A rigid sphere is placed at the center of the box. The velocity on the sphere surface is set to zero. (B) Error norm vs. mesh size for stationary rigid sphere in a linear shear flow

on the shear plane. Dupont et al.(13) demonstrated that an elastic capsule with a prolate spheroid rest shape, whose axis of symmetry is oriented off the shear plane, will exhibit a unique final dynamical motion for all initial orientations. Depending on the capillary number, they observed three final dynamical states: (i) rolling for lower capillary numbers, (ii) wobbling in which the capsule processes around the vorticity axis as the capillary number is increased, and (iii) a swinging-oscillating motion in which the long axis of the capsule oscillates around the shear plane with decreasing amplitude of oscillation as the capillary number increases, resulting in an in-plane swinging motion at high capillary numbers. Wang et al.(14) investigated the off-plane motion of oblate and prolate capsules and concluded that the final dynamical state could depend on the initial inclination angle. A recent study of RBCs in shear flow (15) has demonstrated that RBCs first tumble, then roll, transit to a rolling and tumbling stomatocyte, and finally attain polylobed shapes with increasing shear rate when the viscosity contrast between cytosol and blood plasma is large enough. Minetti et al. (16) give an exhaustive description of the dynamics under a shear flow of a large number of RBCs in a dilute regime is proposed. They identify which of the characteristic parameters of motion and of the transition thresholds depend on flow stress only or also on suspending fluid viscosity.

Similarly, Cordosco and Bagchi (17) studied the off-plane motion of oblate, prolate, and biconcave capsules. Unlike Dupont et al.(13) and Wang et al.(14), they included membrane bending stiffness in their formulation and considered a spatially uniform spontaneous curvature in the case of biconcave capsules. They found that rolling was the dominant mode in the physiologically relevant viscosity ratio case (i.e., 5), tank-treading or wobbling mode at $\lambda < 1$, and an intermittent regime at low capillary numbers and low viscosity ratios, where the dynamics are dependent on the initial orientation. It is noteworthy that Bitbol(18) and Dupire et al.(13) experimentally observed rolling dynamics in a dextran solution where the viscosity ratio was less than unity. The discrepancy between simulation and experiment may result from the use of a spatially uniform spontaneous curvature that corresponds to a biconcave shape. It is important to note that to model the RBC membrane correctly, an assumption of the spontaneous shape has to be made, and finding the appropriate shape has been a challenge for both theoreticians and experimentalists. Sinha et al.(8) investigate the cell dynamics' dependence on the membrane's spontaneous curvature. They found that an oblate spheroidal spontaneous curvature maintains the dimple of the RBC during tank-treading dynamics and exhibits off-shear-plane, tumbling consistent with the experimental observations of Dupire et al.(19). For a complex structure such as an RBC membrane, it is possible that the natural shape for shear elasticity may differ from that for bending elasticity so the overall natural shape of an element results from the balance of bending and shear forces.

There have been endeavors to comprehend the impact of spontaneous shape on the ultimate dynamics of RBCs. Peng et al.(20) conducted a study on the influence of non-biconcave spontaneous shape on RBC dynamics and concluded that in order for an RBC to maintain its biconcave shape during tank-treading, as noted by Dupire et al.(19), the spontaneous curvature must be non-biconcave. In instances where a biconcave spontaneous curvature was employed, tank-treading could not be achieved without significantly perturbing the initial shape. Additionally, Cordosco et al.(21) explored non-biconcave spontaneous shapes and ascertained that the spontaneous shape has a significant impact on cell dynamics, depending on the viscosity ratio. They observed that the dimple in the RBC remained intact for both biconcave and oblate spontaneous shapes. However, it should be noted that in both works, Peng et al.(20) and Cordasco et al.(21), non-biconcave spontaneous curvatures were investigated under the imposition of spatially uniform spontaneous curvature, denoted as $c_0$. It is worth highlighting that RBC membranes differ from model lipid bilayers in that they possess embedded proteins with an underlying spectrin cytoskeleton and an asymmetric bilayer leaflet composition, all of which modify $c_0$, with proteins in particular, having been demonstrated to preferentially bind via curvature-sensing mechanisms. Hence, it can be argued that $c_0$ would be spatially inhomogeneous. Recently, using two different simulation techniques, Mauer et al. (22) construct a state diagram of RBC shapes and dynamics in shear flow as a function of shear rate and viscosity contrast (19, 23). Their studies suggest that a nearly spherical stress-free shape best reproduces experimental results for the tumbling-to-tank-treading transition at low viscosity contrasts. Reichel et al.(24) combined simulation and experimental investigation of RBC shapes and dynamics in microchannels to provide a consistent RBC state diagram and illustrate the complexity of RBC behavior in the microflow. The RBC model employs a stress-free shape of the elastic spring network, corresponding to a spheroidal shape with a reduced volume of 0.96. Their simulation results agree well with experimental observations, allow the characterization of RBC variability in shear elasticity, and permit us to make a significant step toward quantitative measurements of RBC mechanical properties.

In our work, to investigate the effects of membrane curvature on the RBC dynamics, two types of homogeneous normal RBCs suspension are simulated: one with spontaneous bending curvature being biconcave discoid, while another being oblate spheroid. Various vascular geometries are considered, including a slit (Fig.S5), a straight cylindrical tube (Fig. S6), and a curved (serpentine) channel (Fig. S7). The simulation snapshots tell that RBC rest shape has a nontrivial impact on its near-wall dynamics: RBCs with biconcave discoid rest shape tend to "rolling", while RBCs with oblate spheroid rest shape perform "tank-treading", consistent with the findings in a prior numerical investigation by Sinha et al. (8). In spite of the change in orientational dynamics when the spontaneous shape is changed, no substantial change is observed in the number density distribution for normal RBCs.

### Comparison of Segregation in the Straight and Curved Channel

To quantify the segregation between normal and aberrant RBCs, the collective cellular component distribution is measured by the root-mean-square (RMS) distribution of each type of cell from the cylindrical centerline:

$$s = \left\langle {r_{cm}}^2 \right\rangle^{1/2} / a \qquad [9]$$

where $r_{cm}$ is the center-of-mass position of a cell in the radial direction and angle brackets denote averaging over the cells in the system. For the curved channel, $r_{cm}$ is the distance from the cell center-of-mass position to the serpentine channel centerline, as defined previously. Thus a larger $s$ means this type of cell is generally closer to the walls. The time evolution of $s$ for normal and aberrant RBCs from SCD RBC suspensions within both straight cylindrical tubes and serpentine channels are presented in Fig.S8. It is found that at the beginning of the simulation, $s$ increases significantly for aberrant RBCs until reaching a plateau at $s_{aberrant} \approx 3.8$ and $s_{normal} \approx 3.0$. A similar trend is observed in both straight and serpentine channels, suggesting a segregation behavior between normal and aberrant RBCs in SCD RBC suspension. Furthermore, we also note that after the simulation start, $s$ for aberrant cells in the serpentine channel grows faster than that in the straight channel, indicating that cellular segregation occurs more rapidly in the curved channel.

### Simulation snapshots of Cell Distributions in Straight Cylindrical Tube

The simulation snapshots showing the segregation phenomenon between normal and aberrant cells in the straight cylindrical tube are given in Fig. S9.

### RBC-induced Local Wall Shear Stress Fluctuation on Curved Channel Surface

Aberrant RBCs margination provokes local wall shear stress fluctuation on serpentine channel surfaces are shown in Fig. S10, S11, and S12.

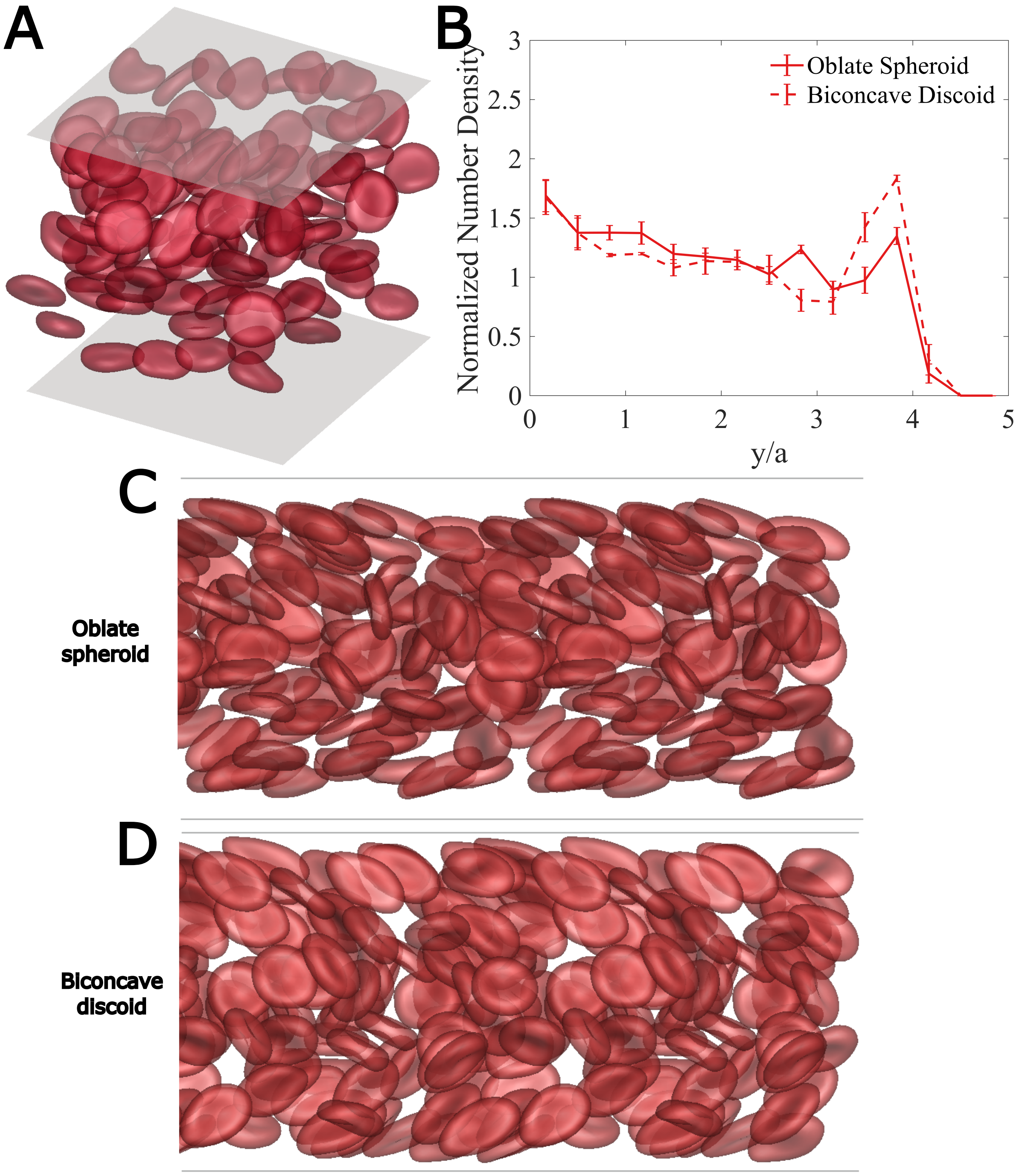


**Fig. S5.** (Top) (A) Simulation diagram of homogeneous normal RBC suspension between the slit under pressure-driven flow. ($Re_p$ = 0.1, Hematocrit = 0.15)(B) Steady-state wall-normal direction cell number density profile. Note here $y/a = 0$ denotes the slit center and $y/a = 5$ close to the wall. (Bottom) The side view of RBC suspension with bending spontaneous curvature being (C) oblate spheroid and (D) biconcave discoid.

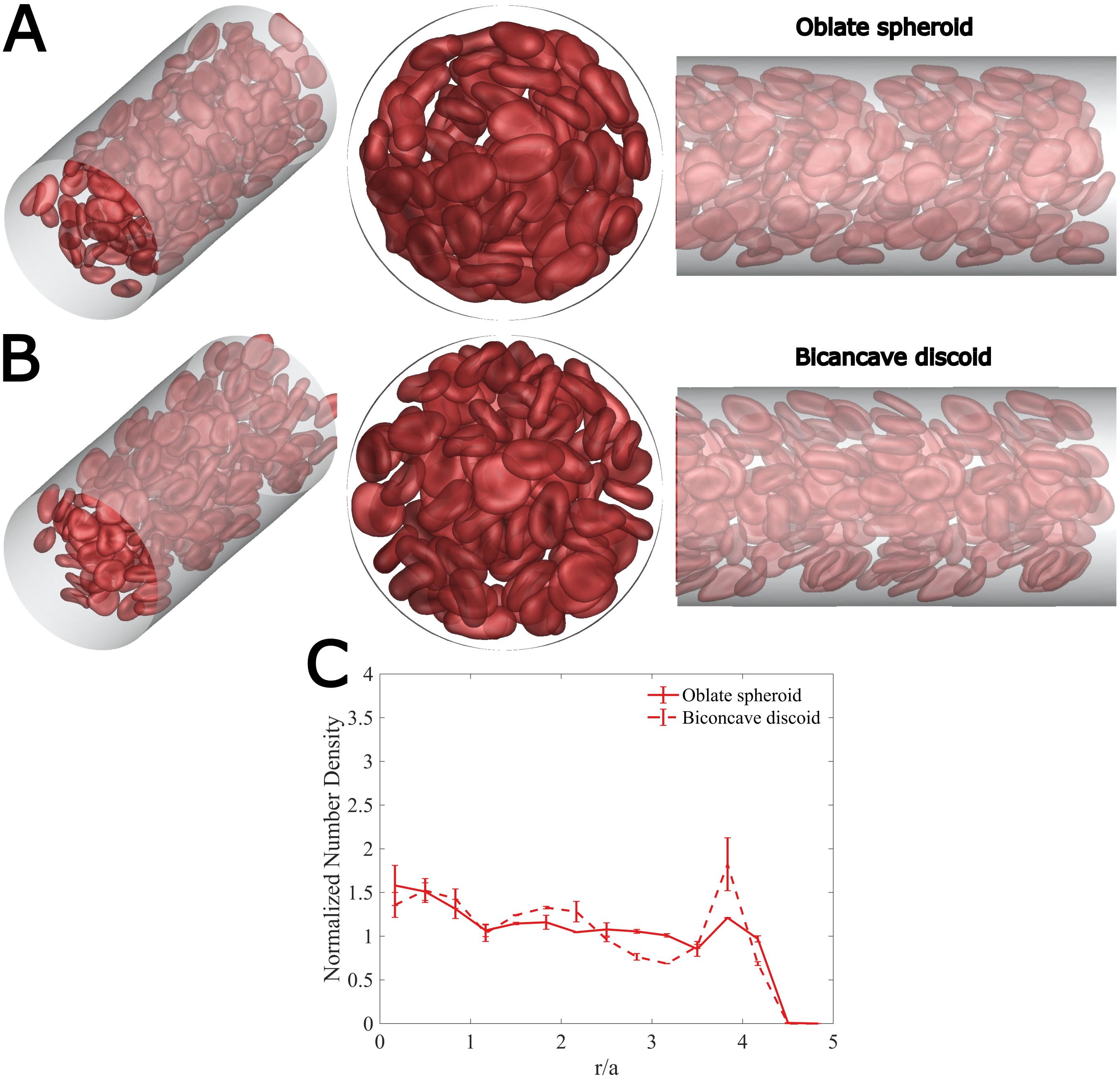


**Fig. S6.** (Top)Simulation snapshots for homogeneous normal RBC suspension within the straight cylindrical tube with spontaneous bending curvature being (A) oblate spheroid and (B) biconcave discoid. ($Re_p$=0.1, Hematocrit = 0.20). (Bottom) (C) Steady-state radial cell number density profile for two types of RBC suspensions.

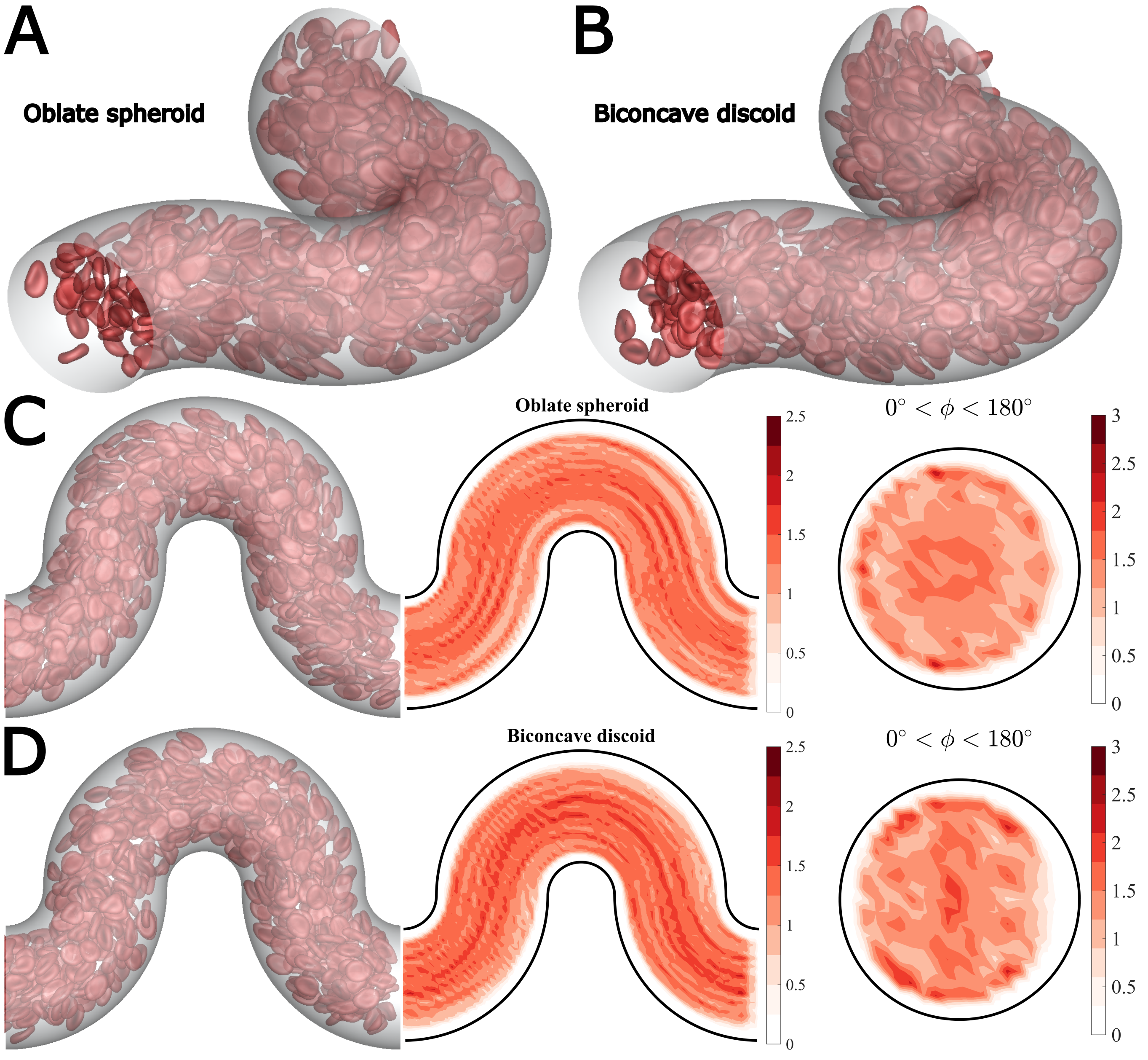


**Fig. S7.** (Top) Simulation snapshots for normal RBC suspension in the serpentine channel with spontaneous curvature being (A) oblate spheroid and (B) biconcave discoid. (Bottom) Center-plane and cross-sectional cell number density distribution for RBC suspension with spontaneous curvature being (C) oblate spheroid and (D) biconcave discoid.

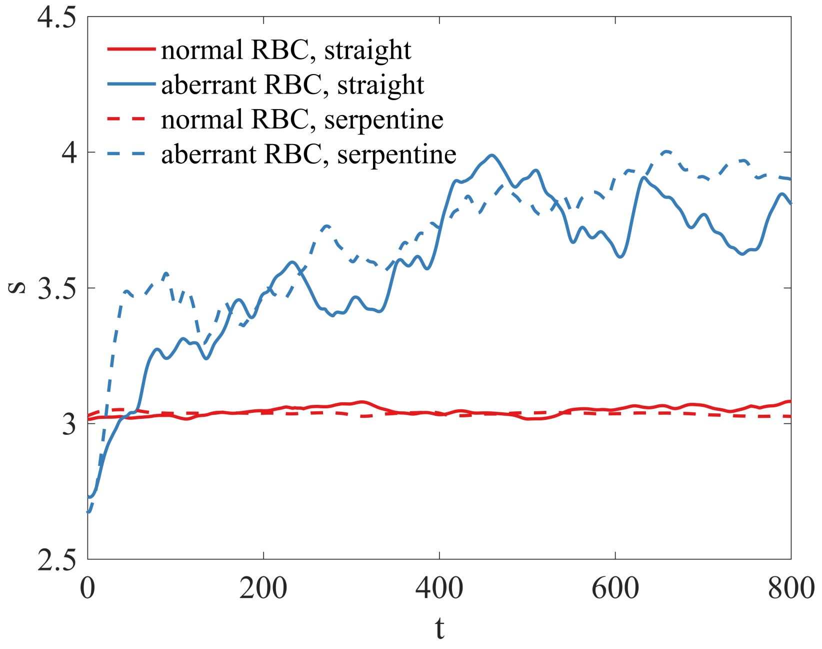


**Fig. S8.** Time evolution of $s$ for each cellular component (red, normal RBCs; blue, aberrant RBCs) in SCD RBC suspension within the straight tube (solid line) and serpentine channel (dash line). $s = \left\langle r_{\mathrm{cm}}^2 \right\rangle^{1/2} / a$. Note that for the straight tube case, $r_{cm}$ is the radial distance from the cell center of mass position to the cylindrical centerline, while for the serpentine channel, $r_{cm}$ is the radial distance to the curved channel centerline. Both the cylindrical channel and curved channel have a tube radius of $20\mu m$.

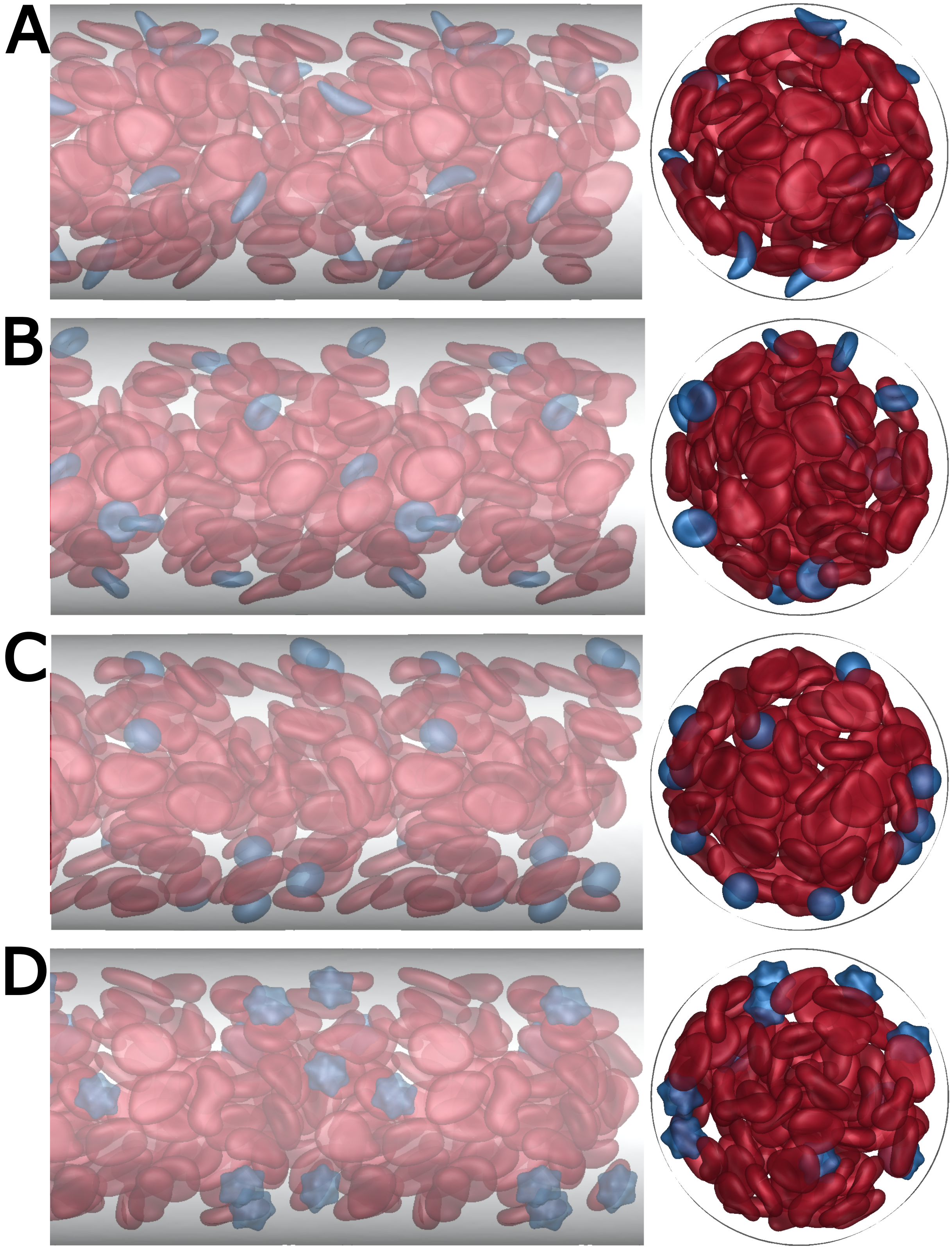


**Fig. S9.** Simulation snapshots (left: side view; right: top view) for (A) SCD, (B) IDA, (C) spherocytosis, and, (D) COVID-19 RBC suspension in the cylindrical straight channel.

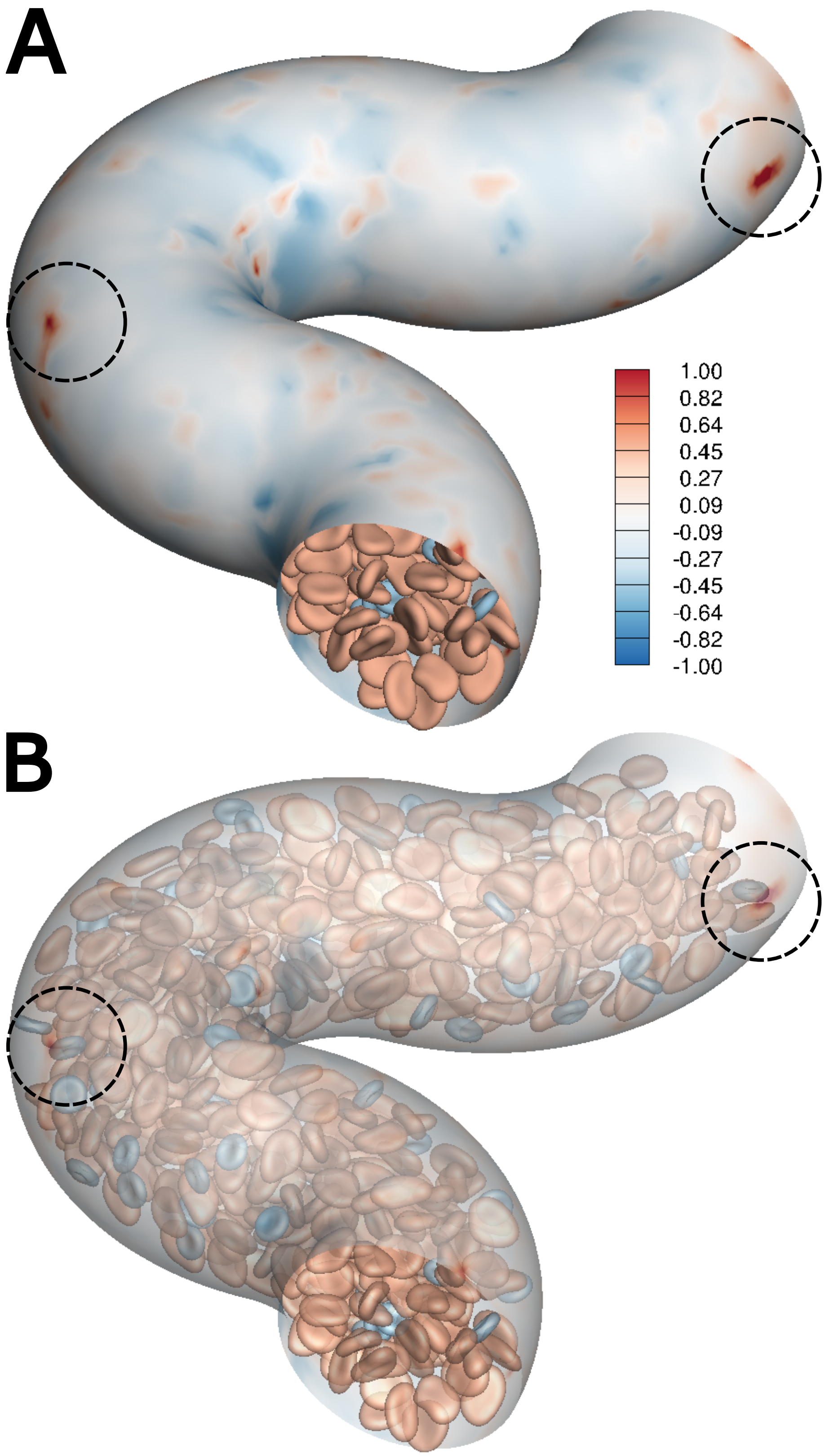


**Fig. S10.** (A) Simulation snapshots and (B) corresponding transparent views of additional wall shear stress $\hat{\tau}_w$ induced by the presence of the cells in suspensions of normal RBCs with iron deficiency RBCs within the serpentine channel. The color on the serpentine vascular surface denotes the RBC-induced wall shear stress strength $\hat{\tau}_w$.

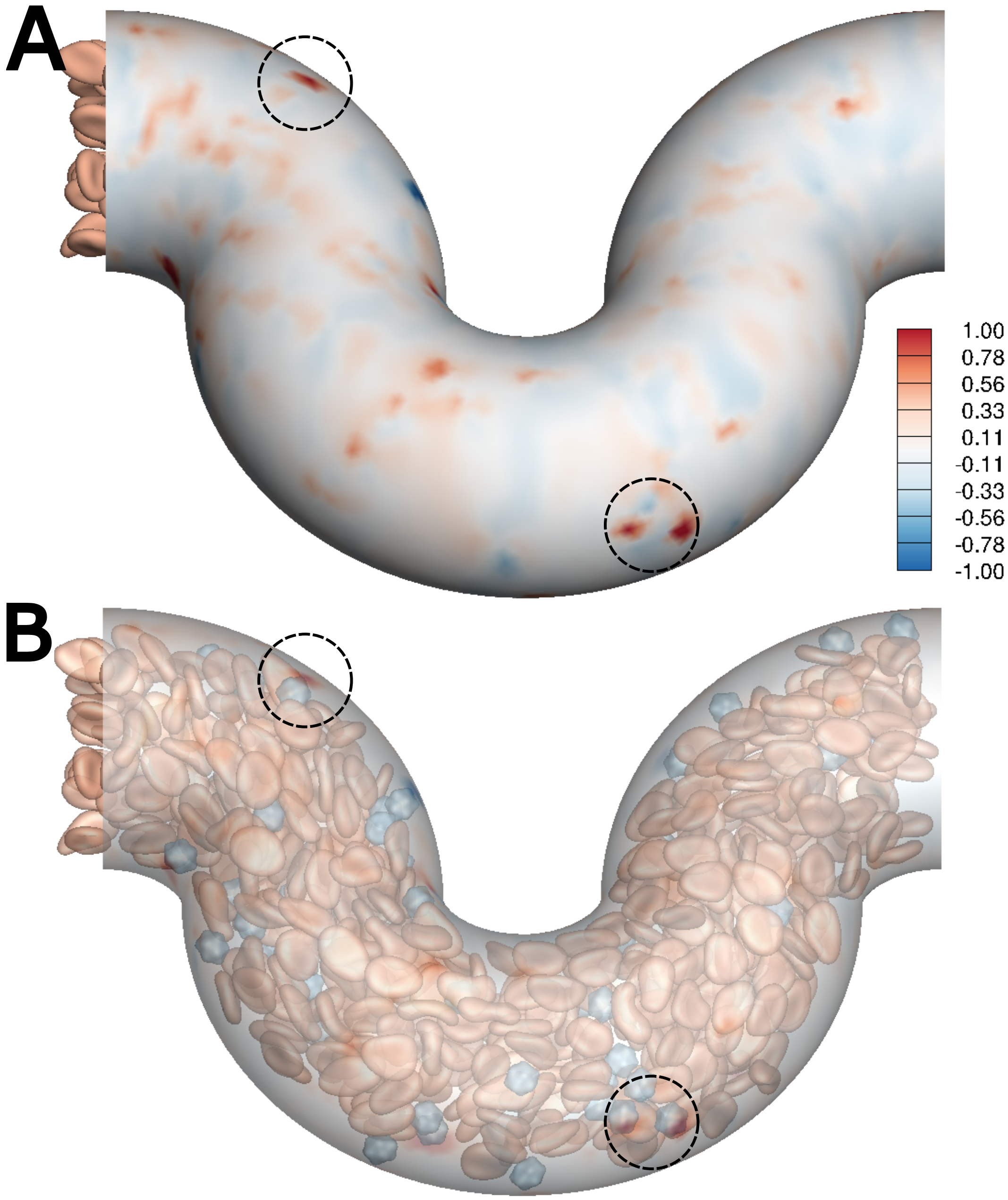


**Fig. S11.** (A) Simulation snapshots and (B) corresponding transparent views of additional wall shear stress $\hat{\tau}_w$ induced by the presence of the cells in suspensions of normal RBCs with sphero-echinocytes within the serpentine channel. The color on the serpentine vascular surface denotes the RBC-induced wall shear stress strength $\hat{\tau}_w$.

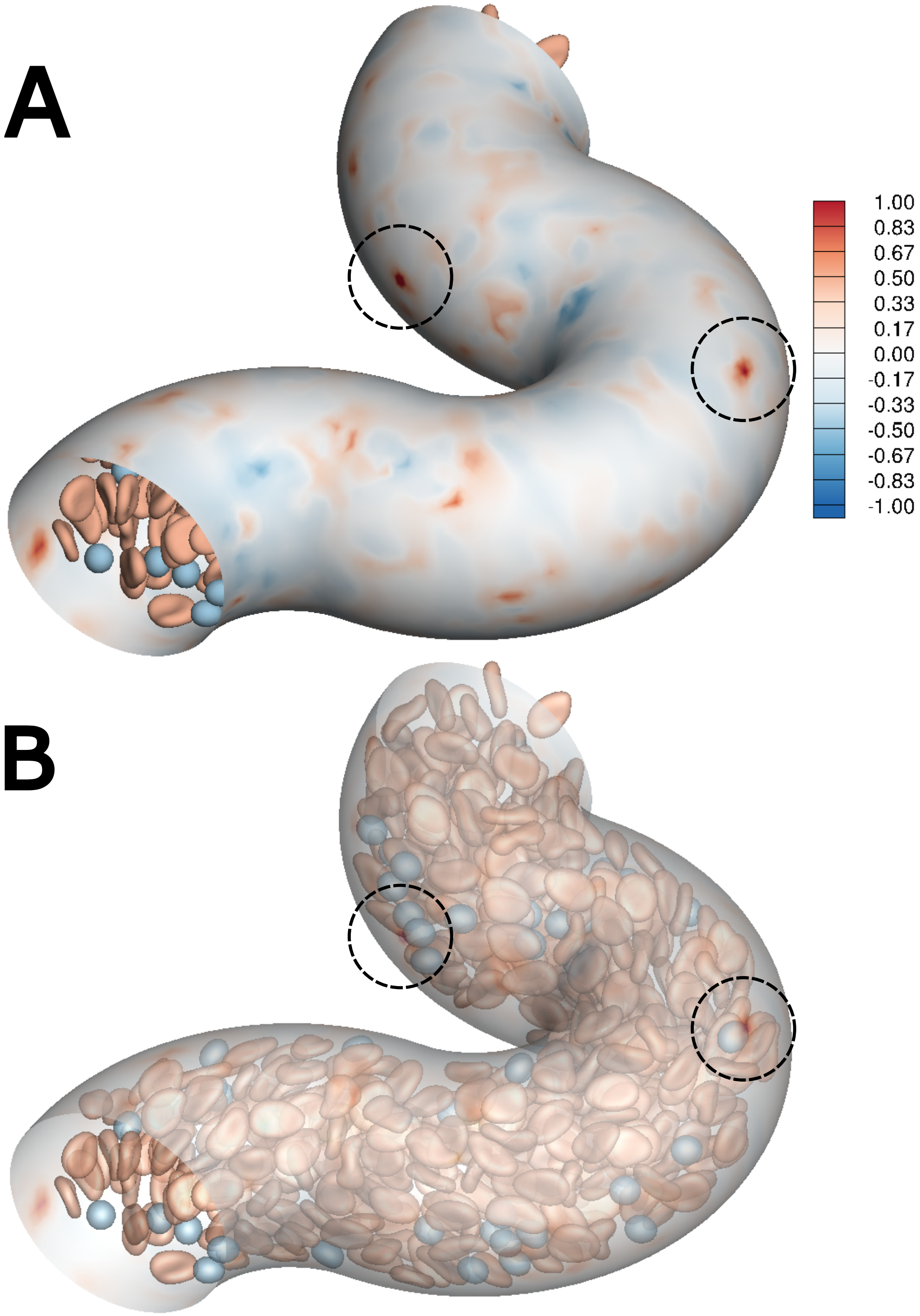


**Fig. S12.** (A) Simulation snapshots and (B) corresponding transparent views of additional wall shear stress $\hat{\tau}_w$ induced by the presence of the cells in suspensions of normal RBCs with spherocytes within the serpentine channel. The color on the serpentine vascular surface denotes the RBC-induced wall shear stress strength $\hat{\tau}_w$.

**Movie S1. SCD RBC suspension in Straight Cylindrical Tube**

**Movie S2. IDA RBC suspension in Straight Cylindrical Tube**

**Movie S3. COVID-19 RBC suspension in Straight Cylindrical Tube**

**Movie S4. Spherocytosis RBC suspension in Straight Cylindrical Tube**

**Movie S5. SCD RBC suspension in Curved Channel**

**Movie S6. IDA RBC suspension in Curved Channel**

**Movie S7. COVID-19 RBC suspension in Curved Channel**

**Movie S8. Spherocytosis RBC suspension in Curved Channel**

### References


1. R Mittal, et al., A versatile sharp interface immersed boundary method for incompressible flows with complex boundaries. *J. computational physics* **227**, 4825–4852 (2008).
2. P Balogh, P Bagchi, A computational approach to modeling cellular-scale blood flow in complex geometry. *J. computational physics* **334**, 280–307 (2017).
3. C Geuzaine, JF Remacle, Gmsh: A 3-d finite element mesh generator with built-in pre-and post-processing facilities. *Int. journal for numerical methods engineering* **79**, 1309–1331 (2009).
4. KC Ong, MC Lai, Y Seol, An immersed boundary projection method for incompressible interface simulations in 3d flows. *J. Comput. Phys.* **430**, 110090 (2021).
5. PB Canham, The minimum energy of bending as a possible explanation of the biconcave shape of the human red blood cell. *J. theoretical biology* **26**, 61–81 (1970).
6. W Helfrich, Elastic properties of lipid bilayers: theory and possible experiments. *Zeitschrift für Naturforschung c* **28**, 693–703 (1973).
7. R Skalak, A Tozeren, R Zarda, S Chien, Strain energy function of red blood cell membranes. *Biophys. journal* **13**, 245–264 (1973).
8. K Sinha, MD Graham, Dynamics of a single red blood cell in simple shear flow. *Phys. Rev. E* **92**, 042710 (2015).
9. G Tryggvason, et al., A front-tracking method for the computations of multiphase flow. *J. computational physics* **169**, 708–759 (2001).
10. JB Freund, J Vermot, The wall-stress footprint of blood cells flowing in microvessels. *Biophys. J.* **106**, 752–762 (2014).
11. P Balogh, P Bagchi, Three-dimensional distribution of wall shear stress and its gradient in red cell-resolved computational modeling of blood flow in in vivo-like microvascular networks. *Physiol. reports* **7**, e14067 (2019).
12. DV Le, ST Wong, A front-tracking method with catmull–clark subdivision surfaces for studying liquid capsules enclosed by thin shells in shear flow. *J. Comput. Phys.* **230**, 3538–3555 (2011).
13. C Dupont, AV Salsac, D Barthès-Biesel, Off-plane motion of a prolate capsule in shear flow. *J. Fluid Mech.* **721**, 180–198 (2013).
14. Z Wang, Y Sui, PD Spelt, W Wang, Three-dimensional dynamics of oblate and prolate capsules in shear flow. *Phys. Rev. E* **88**, 053021 (2013).
15. L Lanotte, et al., Red cells' dynamic morphologies govern blood shear thinning under microcirculatory flow conditions. *Proc. Natl. Acad. Sci.* **113**, 13289–13294 (2016).
16. C Minetti, V Audemar, T Podgorski, G Coupier, Dynamics of a large population of red blood cells under shear flow. *J. fluid mechanics* **864**, 408–448 (2019).
17. D Cordasco, P Bagchi, Orbital drift of capsules and red blood cells in shear flow. *Phys. Fluids* **25**, 091902 (2013).
18. M Bitbol, Red blood cell orientation in orbit c= 0. *Biophys. journal* **49**, 1055–1068 (1986).
19. J Dupire, M Socol, A Viallat, Full dynamics of a red blood cell in shear flow. *Proc. Natl. Acad. Sci.* **109**, 20808–20813 (2012).
20. Z Peng, A Mashayekh, Q Zhu, Erythrocyte responses in low-shear-rate flows: effects of non-biconcave stress-free state in the cytoskeleton. *J. fluid mechanics* **742**, 96–118 (2014).
21. D Cordasco, A Yazdani, P Bagchi, Comparison of erythrocyte dynamics in shear flow under different stress-free configurations. *Phys. Fluids* **26**, 041902 (2014).
22. J Mauer, et al., Flow-induced transitions of red blood cell shapes under shear. *Phys. review letters* **121**, 118103 (2018).
23. M Abkarian, M Faivre, A Viallat, Swinging of red blood cells under shear flow. *Phys. review letters* **98**, 188302 (2007).
24. F Reichel, et al., High-throughput microfluidic characterization of erythrocyte shapes and mechanical variability. *Biophys. journal* **117**, 14–24 (2019).
